# Assessing The Repeatability of Multi-Frequency Multi-Layer Brain Network Topologies Across Alternative Researcher’s Choice Paths

**DOI:** 10.1101/2021.10.10.463799

**Authors:** Stavros I. Dimitriadis

## Abstract

There is a growing interest in the neuroscience community on the advantages of multilayer functional brain networks. Researchers usually treated different frequencies separately at distinct functional brain networks. However, there is strong evidence that these networks share complementary information while their interdependencies could reveal novel findings. For this purpose, neuroscientists adopt multilayer networks, which can be described mathematically as an extension of trivial single-layer networks. Multilayer networks have become popular in neuroscience due to their advantage to integrate different sources of information. Here, we will focus on the multi-frequency multilayer functional connectivity analysis on resting-state fMRI recordings. However, constructing a multilayer network depends on selecting multiple pre-processing steps that can affect the final network topology. Here, I analyzed the fMRI dataset from a single human performing scanning over a period of 18 months (84 scans in total), and the second dataset of 25 subjects with 3 repeat scans. I focused on assessing the reproducibility of multi-frequency multilayer topologies exploring the effect of two filtering methods for extracting frequencies from BOLD activity, three connectivity estimators, with or without a topological filtering scheme, and two spatial scales. Finally, I untangled specific combinations of researchers’ choices that yield consistently brain networks with repeatable topologies, giving us the chance to recommend best practices over consistent topologies.

## 1. INTRODUCTION

New developments in multimodal neuroimaging provide novel directions for measuring structural (anatomical) and functional connectivity (Tulay et al. 2019). These novel developments boost the emergence of brain connectivity (Sporns 2011). An association exists between behavior and cognition and the brain’s large-scale neuronal activity across spatially distributed brain areas (Mišić and Sporns 2016; Alderson et al. 2020). Structural and functional connections between spatially distributed brain areas are recognized as the key element of cognitive functions and behavioral repertoire (Smith et al. 2015; Mišić and Sporns 2016). Recent innovations in non-invasive imaging techniques have assisted neuroscientists to construct comprehensive anatomical maps between neuronal elements (Sporns 2014), and the simultaneous acquisition of dynamic functional brain activity patterns (Loued-Khenissi et al. 2018). The brain is a complex system that can be described as a network (graph) where brain areas are the nodes and their links represent the functional and structural interactions between brain areas (Stam, 2014). The modeling of the brain as a network with any type of neuroimaging modality opens new avenues of graph-theoretic approaches and methods in multiple research directions (Bassett and Sporns 2017).

Network or graph theory has been successfully applied to any neuroimaging modality across many, for example, functional magnetic resonance imaging (fMRI) (Lv et al. 2018), magnetoencephalography (MEG) (Pusil et al. 2019), electroencephalography (EEG) (Maturana-Candelas et al. 2019), diffusion magnetic resonance imaging (dMRI) (Dimitriadis et al., 2017b; Messaritaki et al. 2019), and structural covariance (Carmon et al. 2020). Network theory enables us to simultaneously characterize the spatial organization (network topology) and the strength of any type (either structural or functional) connections (Bertolero and Bassett 2020). Various network metrics that describe nodal (local) and global network characteristics like segregation, integration, (Rubinov and Sporns 2010), and modularity (Sporns and Betzel 2016) have demonstrated their ability to describe quantitatively brain networks in various scientific pathways like in brain diseases (Crossley et al. 2014) and to discriminate brain states while subjects performing cognitive tasks (Braun et al. 2015).

The success of complex network theory in uncovering the key mechanisms of the human brain organization is limited by the use of single-layer brain networks that capture only a single type of interaction (De Domenico 2017). Functional neuroimaging modalities like MEG, EEG, and fMRI can capture brain activity across multiple frequencies and experimental time and it is important to explore the full spectrum (De Domenico et al. 2016; Dimitriadis et al. 2018a; Naro et al. 2021). In contrast, structural neuroimaging modalities such as diffusion-weighted imaging (DWI) measure the presence and strength of physical, and anatomical connections between the various brain areas (Garcés et al. 2016). The necessity of taking advantage of the increasing large multimodal open dataset repositories (Eickhoff et al. 2016) leads to the search for a new type of complex network that can encapsulate functional interactions across multiple frequency scales (multi-frequency case), across experimental time (multi-layer dynamic case), and across modalities (multi-modal case). However, trivial complex networks cannot provide neuroscientists with a mathematical framework to model all the existing interactions across frequencies, time, and modalities.

To present a solution to all the aforementioned challenges, recent research articles in network neuroscience have started to investigate the employment of multilayer networks. A multilayer network enables the integration of information from single-layer networks with the incorporation of interconnected layers that connect these networks (Joseph et al. 2014). Into these current trends, recent research directions in network neuroscience have begun to investigate the employment of multilayer networks to model the multiplex associations that traditional networks are not suited to capture (Boccaletti et al. 2014; Van Mieghem 2016; Muldoon and Bassett 2016; De Domenico 2017). Last years, multilayer networks have been introduced to the network neuroscience field (Brookes et al. 2016; Tewarie et al. 2016; Yu et al. 2017; Dimitriadis et al. 2018a; Buldú and Porter 2018), where different layers correspond to different frequency-dependent functional interactions or to networks derived from different modalities or to specific snapshots of a dynamic functional connectivity network (Battiston et al. 2017).

In the present study, I will focus on multi-frequency multilayer networks, and it is important to mention an important aspect of the construction of this type of multilayer network. Previous neuroimaging studies reported important findings based on multi-frequency multilayer networks. However, the inter-layer connections between frequency-dependent layers were defined as pseudo-links between homologous brain areas between the layers. This practically means that the inter-layer networks involve artificial links that interconnect each node with its representation across layers (Yu et al. 2017; Guillon et al. 2017). However, a true multi-frequency multilayer network should involve also inter-frequency layers that tabulate the cross-frequency interactions between the studying frequencies (Brookes et al. 2016; De Domenico et al. 2016; Tewarie et al. 2016; Dimitriadis et al. 2018a; Williamson et al. 2021).

The spectral features of the resting-state BOLD fMRI (rs-fMRI) multi-ROI signal are of high significant interest (Kalcher et al. 2014). Scientists discovered an alignment between the frequency spectrum within the bandwidth 0 - 0.25 Hz with biological brain mechanisms (Hocke et al. 2016 ;Golestani et al. 2015). Specific spectral content has been associated with both vascular and physiological processes (Golestani et al. 2015 ;Hocke et al. 2016 ;(Mark et al. 2015) and also with derived brain-network connectivity measures (Nikolaou et al. 2016). A few studies attempted to decompose resting-state BOLD activity with either wavelet decomposition (Zhang et al. 2016) or adaptive filtering like empirical mode decomposition (EMD) (Yuen et al. 2019). Here, we will adopt both methods to decompose the rs-fMRI multi-ROI time series into its intrinsic brain frequencies in a data-driven manner.

The construction of multi-frequency multilayer networks can assist researchers to understand better neurodegenerative (Guillon et al. 2017) and psychiatric disorders (Gifford et al. 2020) and they can further design proper connectomic biomarkers. However, these multi-layer based connectomic biomarkers should be repeatable and subject-specific.

A tremendous amount of neuroimaging research articles adopted resting-state fMRI to define repeatable connectomic biomarkers for many brain disorders and diseases (Parkes et al. 2020). Research findings on multi-frequency multilayer networks at resting-state fMRI (rs-fMRI) are preliminary (De Domenico et al. 2016). To design repeatable connectomic biomarkers from resting-state fMRI, a prerequisite is the test-retest repeatability of network topologies (Luppi and Stamatakis 2021).

The majority of rs-fMRI studies adopted multilayer networks to model dynamic functional connectivity interactions in many target disease groups attempting to design connectomic biomarkers of clinical importance (dynamic case ; (Braun et al. 2015; Muldoon and Bassett 2016; Gifford et al. 2020; Dimitriadis et al. 2021). Here, I analyze two open test-retest studies: an rs-fMRI dataset from a single human performing scanning with various modalities over a period of 18 months, and an rs-fMRI dataset from 25 subjects with 3 repeat scans. The application of my analytic framework into two independent datasets with two time spans between scans and a high-frequent long-term scanning of a single subject ensures the generalisability of my findings. My main goal is to assess the reproducibility of multi-frequency multilayer network topologies by investigating the effect of potential choices over a) the filtering method for extracting frequencies from BOLD activity (empirical mode decomposition (EMD) (Yuen et al. 2019) versus wavelet decomposition (Zhang et al. 2016), b) the adopted functional connectivity estimator (Pearson’s correlation coefficient, mutual information, and distance correlation), c) the topological layout of the derived functional brain network (fully-weighted network versus a topological filtering scheme with orthogonal minimal spanning trees (OMST) approach) (Dimitriadis et al. 2017c,a), and d) the spatial scale of the functional brain network (the original based on the parcellation scheme versus a downsampled version based on well-known subnetworks). I adopted portrait divergence PDiV (Bagrow and Bollt 2019) as a proper distance metric to quantify the network topology similarity between every pair of scans and across every set of the aforementioned preprocessing steps (2 × 3 × 2 × 2 = 24 distinct pipelines in total).

To ensure that the proposed multi-frequency multilayer network construction pipelines reduce spurious topological differences, I adopted the following criteria: (i) low PDiV statistically supported compared to the rest of the pipelines, (ii) ability to detect smaller within-subject PDiV than between-subjects PDiV in at least 80% of subjects from the second dataset, and (iii) no significant correlation between PDiV and subject motion, in both datasets (Luppi et al., 2021).

The rest of the paper is organized as follows: Section 2 describes the adopted dataset and the proposed preprocessing framework across multiple levels of choices. Section 3 is devoted to the results of the present study and, finally, section 4 discusses our findings giving instructions to the researchers while presenting the limitations of the current study.

## 2. MATERIALS AND METHODS

### A. Test-Retest Datasets

I analyzed two open test-retest datasets to evaluate the repeatability score of the proposed pipelines. The first dataset involves recordings from one subject over a long period and the second one involves recordings from a group of subjects with short-term and long-term retest periods.

Rs-fMRI was performed in 100 scans throughout the data collection period (89 in the production phase), using a multi-band EPI sequence (TR=1.16_ms, TE=30_ms, flip angle=63 degrees (the Ernst angle for the grey matter), voxel size=2.4 × 2.4 × 2_mm, distance factor = 20%, 68 slices, oriented 30 degrees back from AC/PC, 96 × 96 matrix, 230_mm FOV, MB factor=4, 10:00 scan length). After session no.27, the number of slices was changed to 64 due to an update to the multi-band sequence that increased the minimum TR beyond 1.16 for 68 slices. Finally, 84 sessions were included in the analysis due to the low signal-to-noise ratio (SNR) for 16 sessions (see Poldrack et al. 2015 for further details). The dataset included ten 10-min runs of eyes-closed resting-state data and ten 10-min runs of eyes-open resting-state data. Here, I analyzed only the eyes-closed resting-state recordings. This famous dataset is called MyConnectome and one can test-retest the reproducibility over a long period that is absent in other test-retest studies.

The second open dataset involves the recordings from 25 subjects (mean age 30.7 ± 8.8 years, 16 females) with no history of psychiatric or neurological illness. All participants provided written informed consent compensating for their participation while the study has been approved by review boards of the New York University (NYU dataset) within the School of Medicine. The study has been originally described in (Shehzad et al., 2009), and it is freely accessible from the International Neuroimaging Data-Sharing Initiative (INDI) (http://www.nitrc.org/projects/nyu_trt).

For every subject, 3 resting-state scans were acquired. The first scan was conducted as a baseline, while the second and third scans were conducted on average 11 months later than the first scan (range 5 – 16 months). The second and third scans were conducted on the same day 45 min apart. The contract between the second and third scans will be defined hereafter as a short-term dataset, while the contract between the first scan and the second-third scans will be said hereafter as a long-term dataset.

Scans 2 and 3 were conducted in a single scan session, 45 min apart, which took place on average 11 months (range 5–16 months) after scan 1. Each scan was acquired using a 3T Siemens (Allegra) scanner, and consisted of 197 contiguous EPI functional volumes (TR = 2000 ms; TE = 25 ms; flip angle = 90°; 39 axial 163 slices; field of view (FOV) = 192 × 192 mm2; matrix = 64 × 64; acquisition voxel size = 3 × 3 164 × 3 mm3). Participants were instructed to relax and remain still with their eyes open during the scan. For spatial normalization and localization, a high-resolution T1-weighted magnetization prepared gradient echo sequence was also obtained (MPRAGE, TR = 2500 ms; TE = 4.35 ms; TI = 900 ms; flip angle = 8°; 176 slices, FOV = 256 mm).

### B. Functional MRI preprocessing

All fMRI data were preprocessed according to a pipeline developed at Washington University (Poldrack et al. 2015; see section 1 in supp.material). The parcellation procedures lead to 630 parcels (ROIs). The same parcellation has been adapted also for the second dataset.

### C. Construction of Multi-frequency Multilayer Networks

#### Node definition

Every ROI characterized by a specific frequency content is characterized as a distinct node in the multi-frequency multilayer network. In this study, I will decompose every ROI-based brain activity into four basic frequencies (see next section). This practically means that the size of our multilayer network will be: {4 × 630} x {4 × 630} = 2520 × 2520. This multilayer network will tabulate both the within and between frequencies coupling across every pair of ROIs. In multi-frequency multilayer networks, a node is defined as a frequency-dependent brain activity of every ROI.

#### Extracting of brain frequencies

I extract wavelet coefficients for the first four wavelet scales, which correspond to the frequency ranges 0.125□0.25 Hz (Scale 1), 0.06□0.125 Hz (Scale 2), 0.03□0.06 Hz (Scale 3), and 0.015□0.03 Hz (Scale 4) (Zhang et al. 2016). Here, I adopted the maximum overlap discrete wavelet transform (MODWT), selecting the Daubechies family implemented with a wavelet length equal to 6.

Alternatively, I decompose resting-state BOLD activity into the related intrinsic mode functions (IMFs) with the empirical mode decomposition (EMD) (Yuen et al. 2019). I estimated the mean frequency of the Hilbert spectrum across time per brain area and IMF across scans.

I followed both decomposition methods first on the extracted averaged time series per brain area for every scan across the 630 ROIs for Pearson’s Correlation Coefficient (PC) and Mutual Information (MI) estimations, and secondly on the voxel time series within every ROI per scan for Distance Correlation (DC) estimations. Fig.1 illustrates the decomposition with the two adopted methods of mean representative time series across voxels from the first two ROIs as presented in the MyConnectome dataset. Fig.1A, and B are dedicated to EMD and MODWT, respectively. I constructed multi-frequency multilayer networks using the 8 in total time series (2 ROIs x 4 frequency subbands) and adopted the three connectivity estimators. The estimated network topologies for both decomposition methods are shown on the right side of each sub-figure. Blocks of connectivity strength within the 4 time series per ROI are tabulated within the main diagonal. The off-diagonal blocks tabulate connectivity strengths between the two sets of four-time series. Both PC and MI are estimated on the representative time series per ROI (shown in red) derived from averaging the voxel-based time series (shown in blue). In contrast, DC is computed between two sets of voxel-based time series (shown in blue).

**Figure 1.**
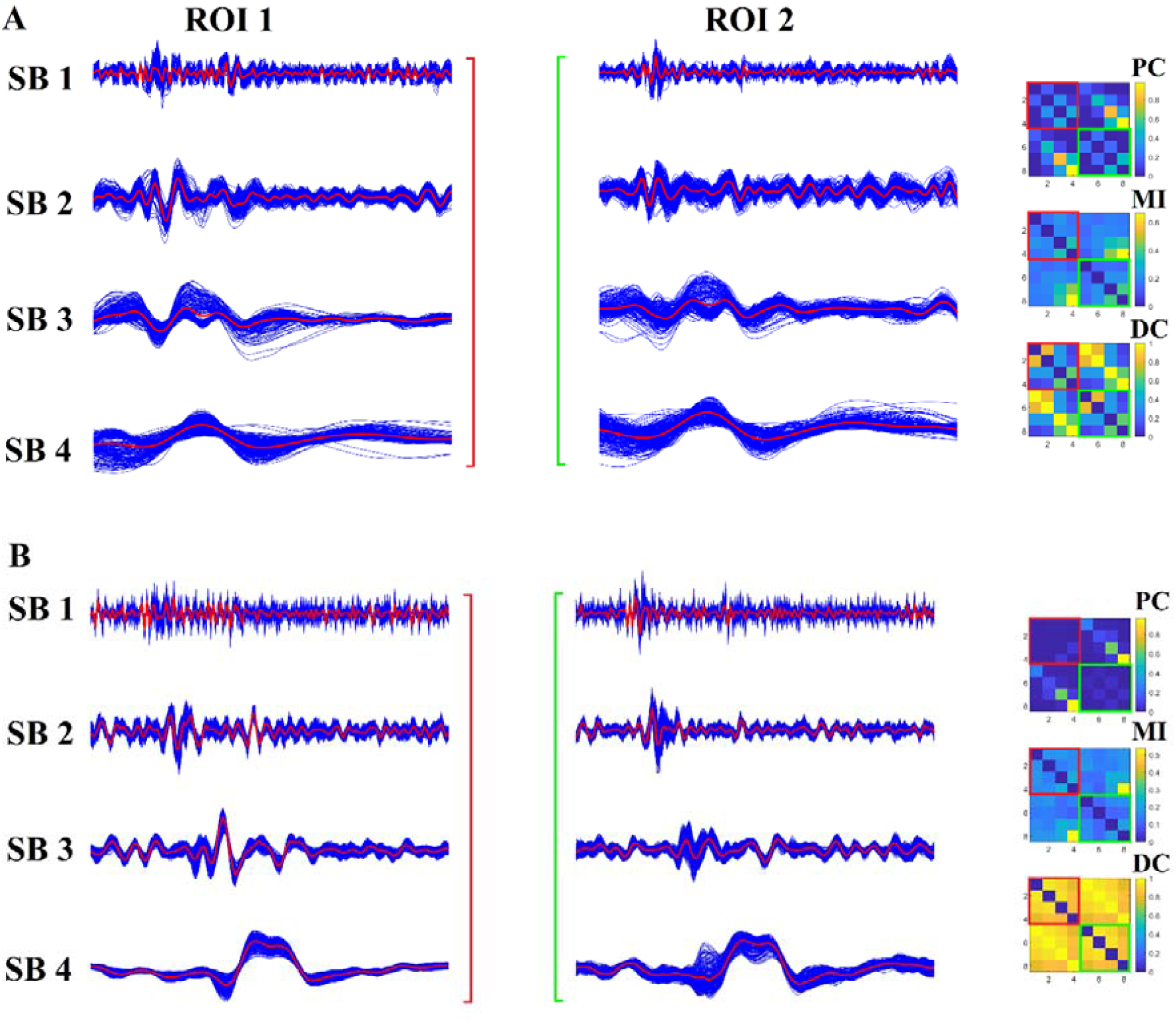
Decomposition of BOLD activity in frequency subbands with EMD (A) and MODWT (B). I showed in blue the voxel-based time series for the first two ROIs of the 1^st^ scan from the MyConnectome project using both decomposition methods. The averaged representative time series is shown in red. Network topologies tabulate the functional connectivity strength across the 8 time series (2 ROIs x 4-time series) with the three adopted connectivity estimators. Blocks within the main diagonal are color-coded to underline the functional interactions between the 4 time series per ROI. The off-diagonal blocks tabulate the functional connectivity strength of the two sets of 4-time series in a pair-wise fashion.

#### Functional Connectivity Estimators

In the present study, I adopted three connectivity estimators that are divided into two groups. The first group involves the Pearson’s Correlation Coefficient (PC), and the Mutual Information (MI). Both estimators can quantify the functional coupling strength between every pair of two frequency-dependent time series derived as the ROI-averaged representative time series. The second group involves the Distance Correlation (DC) metric that can quantify the correlation of two sets of frequency-dependent time series corresponding to the voxel-based time series of two ROIs.

I constructed a multilayer network whose ij^th^ elements are given by the three connectivity estimators with blocks in the main diagonal of size 630 × 630 corresponding to the four within-frequency functional connectivity networks and off-diagonal blocks of size 630 × 630 corresponding to every possible pair of the between-frequency functional connectivity networks (4×3/2=6 in total). The aforementioned procedure was followed for every single scan, filtering method, and connectivity estimator.

For each parcellation, the average denoised BOLD time series across all voxels belonging to a given ROI were extracted. I considered three alternative ways of quantifying the interactions between regional BOLD signal time series.

Below, I defined the mathematical descriptions of the adopted connectivity estimators.

#### Pearson linear correlation

First, I used Pearson correlation, whereby for each pair of nodes *i* and *j*, their functional connectivity strength *FCS*_*ij*_ was given by the Pearson correlation coefficient between the time courses of *i* and *j*, over the full scanning length. I got the absolute Pearson’s correlation values that bound the range of FCS within [0,1].

#### Mutual information (MI)

Second, I also used the mutual information *I*, which quantifies the interdependence between two random variables X and Y, and is defined as the average reduction in uncertainty about X when Y is given (or vice versa, since this quantity is symmetric):

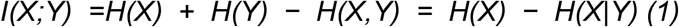

With *H(X)* being the Shannon entropy of a variable X. Unlike Pearson correlation, mutual information considers both linear and nonlinear relationships. I normalized mutual information (MI) values by dividing the maximum value in the matrix-bound within [0,1].

#### Distance Correlation

The multivariate distance correlation (DC) measure was introduced in the literature on functional network construction in general and specifically in rs-fMRI many years ago (Geerligs et al., 2016). However, it was introduced to construct a single-layer functional network based on BOLD activity within a single frequency range. Here, I adopted for the very first time based on the authors’ knowledge distance correlation as a proper functional connectivity estimator for both within and between frequencies coupling in the multi-frequency multilayer functional brain network construction. This new test is based on an <u>unbiased estimator</u> of distance covariance, and the resulting t-test is unbiased for large sample sizes (> 30) (Székely and Rizzo 2013). The combined p-value can be estimated analytically. Here, I adopted distance correlation to estimate the functional connectivity strength between pairs of tuples of voxel-based time series between every pair of ROIs. DC can capture both linear and non-linear associations between two time series.

#### Surrogate Null Models: Statistical Topological Filtering

Since the ground truth of the presence of true functional connections cannot be defined, the construction of surrogate data as a statistical framework is inevitable (Schreiber and Schmitz 2000; Pereda et al. 2005). Surrogate time series must preserve specific properties of the original time series to be useful. These properties are the auto□covariance sequence, stationary cross-correlation, power spectral density, cross power spectral density, and amplitude distribution (Schreiber and Schmitz 2000; Pereda et al. 2005; Zalesky et al. 2014). In the present study, I adopted two basic surrogate data methods: the first one produces surrogate data adopting the notion of the multivariate phase randomization (MVPR) (Prichard and Theiler 1994), and the second is called multivariate autoregressive (MVAR) (Zalesky et al. 2014; Savva et al. 2019).

The MVPR method is first described for generating surrogate time series(Prichard and Theiler 1994). Below, I described briefly the steps of producing the surrogate time series. Let x = [x_1_,x_2_,…,x_n_] denote the BOLD recordings from n=630 parcels each of these time series is composed of 518 time points and X =[X_1_,X_2_,…,X_n_] denote their discrete Fourier transform. Then, I generated a uniformly distributed random phase (φ = [φ_1_,φ_2_,…,φ_τ_]), within the interval [0, 2π] and we further applied to each signal with the following equation: X_k_ = X_k_^eiφ^, k = 1,2,…,n. Practically, this transformation means that in the frequency domain, all our recorded signals are multiplied by the same uniformly random phase (Hindriks et al. 2016). Finally, I estimated the inverse Fourier transform and I got our first surrogate dataset. I repeated the same procedure 1,000 times producing 1,000 surrogate datasets for every scan.

MVAR models produce a set of signals described as a combination of both their own past and also the past of the entire set of signals in the multidimensional set (Prichard and Theiler 1994). The polynomial order p defines the number of past signal values that are considered in the MVAR model. I selected the value of p based on the minimization of the Schwarz Bayesian Criterion (SBC) (Zalesky et al. 2014). Again, a total number of 1,000 randomized copies were created for every time series (Zalesky et al. 2014; Hindriks et al. 2016).

I applied both MVPR and MVAR to the original BOLD time series. To validate the derived surrogate time series with both methods, I adopted the following properties that should be preserved: auto-covariance sequence, stationary cross-correlation, power spectral density, cross power spectral density, and amplitude distribution. For every ROI-based time series and the estimated surrogates, we adapted the absolute value of the Pearson’s correlation coefficient (aPCC) and mutual information (MI) as quality measures of every property. For the auto-covariance sequence, I estimated the aPCC between the auto-covariance sequence of the original time series with every surrogate. I also estimated the cross-correlation between the original time series and every surrogate. For power spectral density (Welch’s power spectral density estimate using *pwelch* MATLAB’s function), I estimated the aPCC between the power spectral densities of the original time series and every surrogate. For cross power spectral density (*cpsd* MATLAB function), I estimated the aPCC between the cross power spectral density of the original time series with itself with the cross power spectral density between the original time series and every surrogate. Finally, the MI was used to quantify the amplitude distributions of the original time series and every surrogate. The whole analysis was repeated per ROI, and the final property values were averaged across the number of surrogates. I repeated this analysis independently per subject, scan, and frequency sub-bands with both filtering methods either EMD or MODWT. For further details, see section 2 in supp.material.

#### Surrogate Null Hypothesis

For every multi-frequency multilayer network, I generated 1,000 surrogate multilayer networks based on both methods. Then, I assigned to every functional connection a *p*-value by estimating the proportion of surrogate connectivity values that were higher than the observed values (Theiler et al. 1992). To correct the effects of multiple comparisons, *p*-values were adjusted using the false discovery rate (FDR) method (Benjamini and Hochberg 1995; Dimitriadis et al. 2015). A threshold of significance *q* was set such that the expected fraction of false positives was restricted to *q* ≤ 0.01 (Dimitriadis et al. 2015; Dimitriadis 2021). The whole procedure was repeated separately across filtering methods, connectivity estimators, and scans. Statistical topological filtering multi-frequency multilayer networks were then fed to our data-driven topological filtering scheme called OMST.

#### Data-driven topological filtering scheme

An important pre-processing step for brain networks is to topologically filter out the backbone of functional links across the whole network. Here, I adopted my data-driven technique called Orthogonal Minimal Spanning Trees (OMST) (Dimitriadis et al. 2017c, a) for the very first time to topologically filter a multilayer network.

OMST (Dimitriadis et al. 2017c, a) is a data-driven approach that optimizes the balance between global efficiency and the global-cost efficiency of the network which is defined as the global efficiency minus the cost. The cost is defined as the ratio of the sum of functional strength of the selected functional links versus the total sum of functional strength of all the pairs of functional links (fully-weighted version of the network). OMST can be described with the following steps : (1) at the first stage, the original MST is extracted consisting of N-1 functional links (where N denotes the total number of nodes) that connect all the nodes while simultaneously minimizing the average wiring cost. MST captures the main net of functional links where the major part of all pairs of shortest paths pass through. Global efficiency and global-cost efficiency are estimated for the 1^st^ MST ; (2) Then, the N-1 functional links were removed from the network, and we searched for the 2^nd^ MST which is orthogonal to the first. We added the 1^st^ and 2^nd^ MST to the network, and we again estimated the global efficiency and the global-cost efficiency ; (3) We repeated the same procedure until a global maximum is detected on the plot of global-cost efficiency versus the total cost (Dimitriadis et al. 2017c, a). The OMST procedure produces sparse functional networks but denser than using only the first MST. Moreover, the OMST method doesn’t impose a-priori selected sparsity level across a cohort, and it produces highly repeatable structural and functional networks compared to alternative topological filtering schemes (Dimitriadis et al. 2017b, 2018b; Messaritaki et al. 2019). Here, I analyzed fully-weighted multilayer networks and topologically filtered multilayer networks with OMST.

#### Network Scales

In the present study, I constructed a multi-frequency multilayer network based on the parcellation scheme provided by the authors of the MyConnectome project (Poldrack et al. 2015). The total number of ROIs as was already aforementioned was 630. Here, I explored the within and between frequency interactions across every pair of ROI for a total of four frequency bands as extracted with MODWT and EMD methods. This practically means that the size of our multilayer network will be equal: {4 × 630} x {4 × 630} = 2520 × 2520. Fig.2 visualizes an example of a multi-frequency multilayer network constructed with the combination of EMD and PC. Fig.2A illustrates the fully-weighted multi-frequency multilayer network while the OMST version of the multilayer network is depicted in Fig.2B. Simultaneously, as many researchers integrated their findings into well-known resting-state networks, I decided to create a subnetwork multilayer network as follow: we computed the mean of pair-wise functional strength between ROIs that comprised each of the following thirteen cognitive networks as provided within the MyConnectome project (Poldrack et al. 2015). These subnetworks are Default Mode Network, Somatomotor, Ventral_Attention, Frontoparietal_1, Frontoparietal 2, Visual_1, Visual_2, Medial_Parietal, Parieto_occipital, Cingulo_opercular, Salience, Dorsal_Attention, and a final subnetwork that includes ROIs that are not classified to the twelve subnetworks. The final size of these subnetworks are equal to : {4 × 13} x {4 × 13} = 52 × 52. An example of a small-scale multilayer network is shown in Fig.3 for the combination of EMD and PD as in Fig.2.

**Figure 2.**
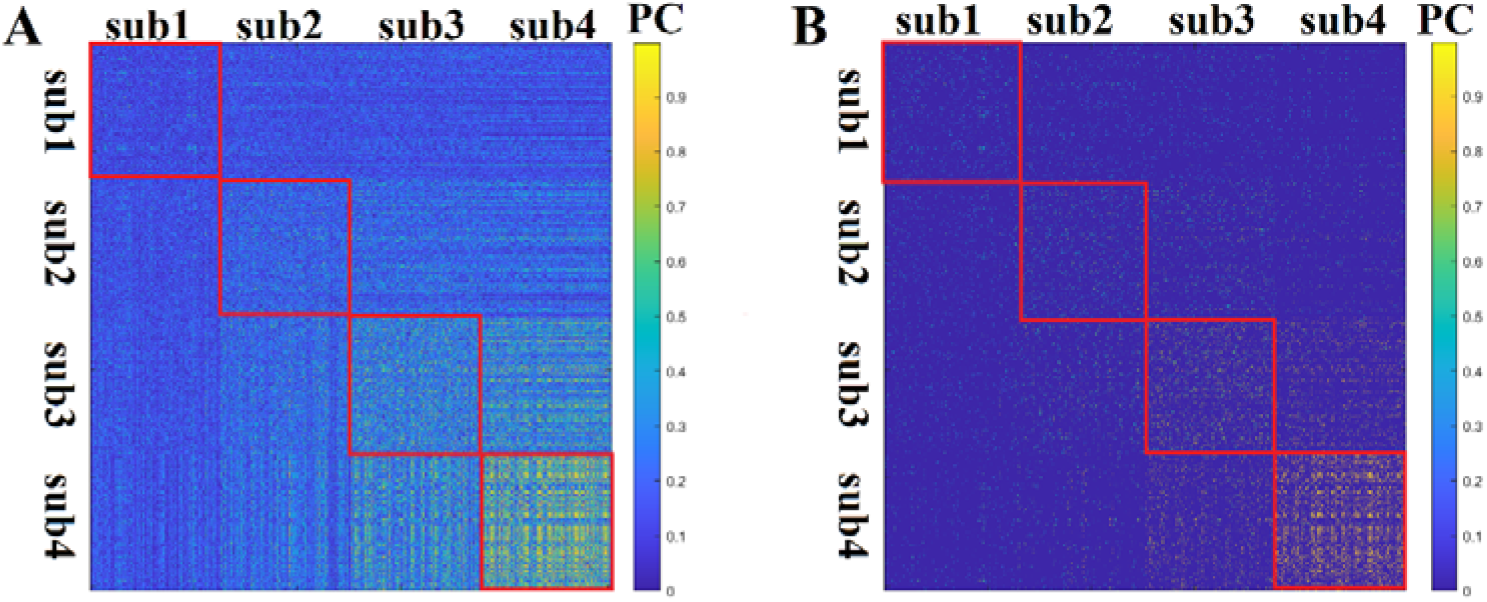
An example of a full-resolution multi-frequency multilayer network from the 1^st^ scan derived from the combination of the EMD filtering technique and PC as a proper functional connectivity estimator. A) A fully-weighted version of the multi-frequency multilayer network B) The OMST version of the multi-frequency multilayer network shown in A In-diagonal red blocks underline the intra-frequency functional networks of size 630×630. Off-diagonal blocks refer to cross-frequency (inter-frequency) functional networks of the same size (**sub-subband**)

**Figure 3.**
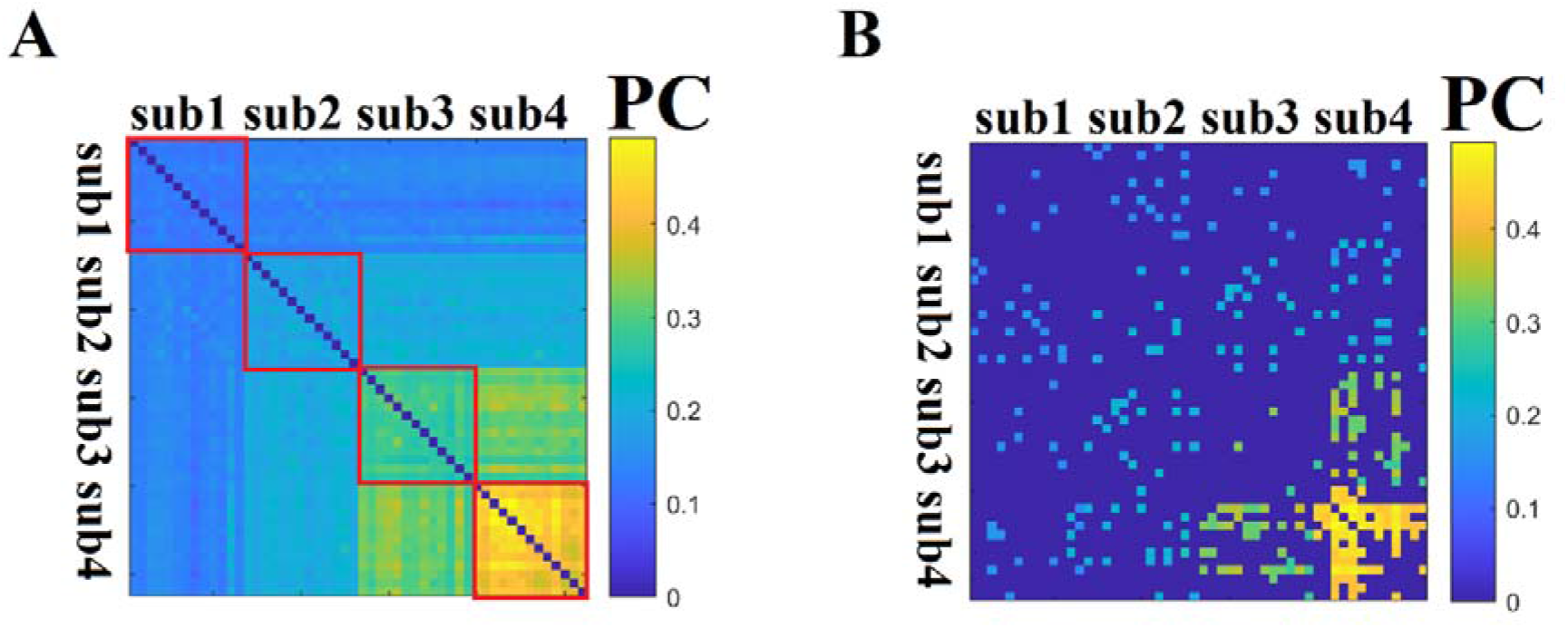
An example of a low-resolution multi-frequency multilayer subnetwork from the 1^st^ scan derived from the combination of the EMD filtering technique and PC as a proper functional connectivity estimator. The size of this subnetwork is 52 × 52. A) The fully-weighted low-resolution multi-frequency multi-layer subnetwork. In-diagonal red blocks underline the intra-frequency functional subnetworks of size 13×13. Off-diagonal blocks refer to cross-frequency (inter-frequency) functional subnetworks of the same size. B) The OMST version of the low-resolution multi-frequency multi-layer subnetwork (**sub - subband**)

### D. Topological Distance as Portrait Divergence

To quantify the difference between network topologies, I used the recently developed Portrait Divergence (PDiV). The Portrait Divergence (PDiV) between two graphs G_1_ and G_2_ is the Jensen-Shannon divergence between their “network portraits”, which encode the distribution of shortest paths of the two networks (Bagrow and Bollt 2019). Specifically, the network portrait is a matrix *B* whose entry *B*_*lk*_, *l* = 0, 1, …, *d* (with *d* being the graph diameter), *k* = 0, 1, …, *N* ™ 1, is the number of nodes having *k* nodes at shortest-path distance *l*. For further details, an interested reader can read the original article describing this method (Bagrow and Bollt 2019).

PDiV considers all the scales of the topology within the networks from motifs to large-scale connectivity patterns and is not restricted to a single network property (Bagrow and Bollt 2019).

For each scan, I obtained one brain network following each of the possible combinations of steps above ((2 decomposition scheme: MODWT/EMD) x (3 connectivity estimators: PC/MI/DC) x (2 topological filtering: full/OMST) x (2 scales: Atlas/Subnetwork) = 24 distinct pipelines in total).

For each pipeline, I then computed the PDiV between multi-frequency multilayer brain network topologies obtained from the single subject at different time points (scans). This procedure resulted in the construction of 24 similarity matrices of size 84×84 (scans x scans) for the first dataset. Every similarity matrix tabulated the PDiV distance of the multilayer brain network topologies related to every scan in a pair-wise fashion. I finally estimated the mean PDiV across every possible pair of 84 scans (84×83/2 = 3486 pairs) to characterize the quality of each of the 24 distinct pipelines. Fig.4 illustrates an example of scan-to-scan pair-wise PDiV distances between every pair of multi-frequency multi-layer networks. The column on the right shows the sum of every row in the distance D matrix called ΣPDiV. This vector of size equal to the number of scans expresses the (dis)similarity of every single-scan multilayer network topology across the rest of the scan-related multilayer network topologies.

**Figure 4.**
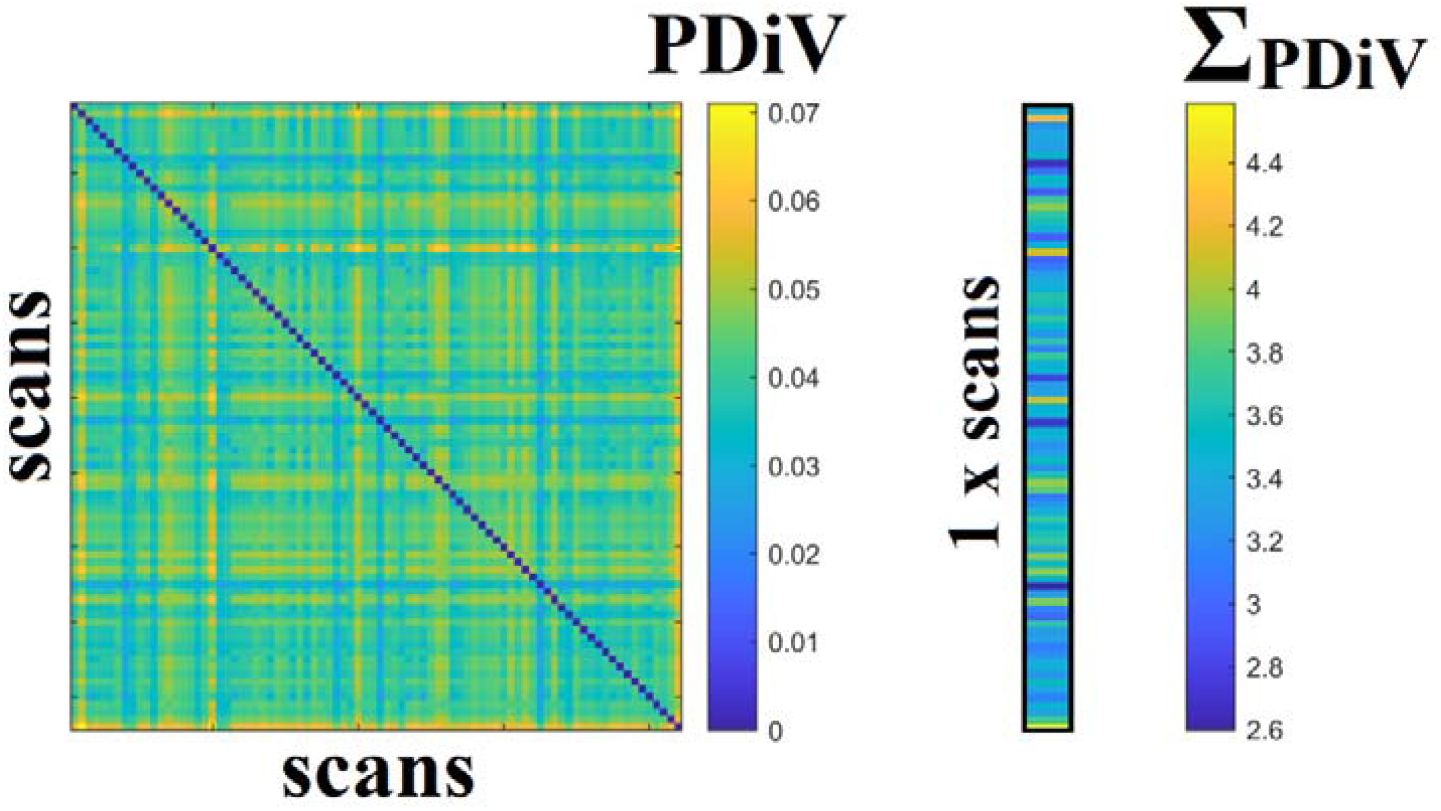
An example of scan-to-scan pairwise topological PDiV distances between pairs of multi-frequency multilayer networks derived from the combination of EMD filtering technique and PC as a proper functional connectivity estimator. The column on the right shows the sum of every row in the distance D matrix called ΣPDiV. The size of this vector is equal to the number of scans.

For the second dataset, I computed the PDiV between multi-frequency multilayer brain network topologies obtained from the 3 repeat scans. Specifically, I quantified the PDiV topological distance between the first and the second – third scans as long-term and the topological distance between the second – third scans as short-term. In the long-term scenario, I got the mean of the two derived PDiV values. The whole analysis is repeated independently per subject and finally, I group-averaged the PDiV values for short-term and long-term across the 24 distinct pipelines.

### E. Criteria for the Evaluation of Multi-Frequency Multilayer Network Construction Pipelines

To ensure that the proposed multi-frequency multilayer network construction pipelines reduce spurious topological differences, I adopted the following criteria across the three cases (the MyConnectome dataset and the two time spans from the NYU dataset). The same criteria were used in our previous study (Luppi et al., 2021):

#### (i) low PDiV statistically supported compared to the rest of pipelines

I set up a criterion of 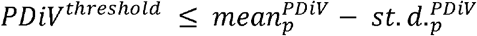 (2) to characterize a pipeline *p* as repeatable. The mean and the standard deviation (st.d.) were estimated across the 24 distinct pipelines independently for the first dataset, and the short-term and long-term scenarios.

#### (ii) ability to detect smaller within-subject PDiV than between-subjects PDiV in at least 80% of subjects from the second dataset

Another criterion that can further evaluate the appropriateness of the pipelines is by direct comparing PDiV values estimated within subjects (WS) (scan 1 vs. scan 2 for subject 1, scan 1 vs. scan 2 for subject 2, etc…) and PDiV values estimated between subjects (BS) (subject 1 vs. subject 2, etc…). The proportion of subjects that showed lower WS^PDiV^ values compared to BS^PDiV^ could be used as an additional quality criterion of the pipeline (Luppi et al., 2021). The rationale behind this criterion is that the individual’s multi-frequency multilayer network topologies between two scans separated in time should differ less than it differs from the multi-frequency multilayer network topology of other individuals. This criterion has been applied to the second NYU dataset for both time spans.

#### (iii) no significant correlation between PDiV and subject motion, in both datasets

As a final criterion, I exclude pipelines whose PDiV value is significantly correlated with differences in subject motion using the mean framewise displacement (Luppi et al., 2021).

### E. Statistical Analysis

#### First Quality Criterion

Scan-averaged PDiV for the MyConnectome study and subject-averaged PDiV for the NYU study were estimated for each of the potential 24 distinct pipelines. To explore the effect of researcher choice at the four levels of preprocessing steps on the repeatability of multi-frequency multilayer topology, I followed an n-way ANOVA (p < 0.05). I performed two three-way ANOVA with repeated measures on three factors (filtering - connectivity estimator - topological layout), one in the atlas and one in the subnetworks space. As an input to the three-way ANOVA for the MyConnectome study, I employed the 84 values produced by the sum of every row of the distance matrix as it was shown in Fig.4. As an input to the three-way ANOVA for the NYU study, I employed the group-averaged PDiV values for both time-sessions. The final p-values of every single pre-processing step and their interactions were adjusted for multiple comparisons in both cases.

#### Second Quality Criterion

I adopted a Wilcoxon Rank-Sum test to evaluate the pipelines proposed by the first PDiV criterion compared to the rest if they showed a higher proportion of subjects with WS^PDiV^ values lower than BS^PDiV^ in both short-term and long-term time spans of the NYU dataset (p < 0.05).

#### Third Quality Criterion

I adopted Pearson’s correlation coefficient to estimate the correlation between the PDiV values and the mean framewise displacement.

#### Evaluation of the best Surrogate Null Model

I also adopted a Wilcoxon Rank-Sum test to compare for every ROI, frequency sub-band, and filtering method, the five properties between the two surrogate null-models (p < 0.05). In the case of the MyConnectome study, I used a vector of 84 values referring to the 84 scans to compare every property, for every ROI between the two surrogate null models. Finally, I reported the percentage of ROIs in both datasets that showed significant differences while I accounted for which surrogate null models demonstrated surrogates with properties closer to the original time series. The findings are reported in section 2 in supp.material.

## RESULTS

### Appropriate Surrogate Model for Statistical Filtering of Multilayer Networks

Produced surrogate time series must preserve specific properties of the original time series in order to be useful. Surrogate BOLD time series produced by the MVPR model preserved better the adopted properties compared to the surrogates produced by the MVAR in a large percentage of ROIs across the two filtering methods and the four subbands (see section 2 and STables 1 – 3 in supp.material).

### Characteristic Intrinsic Frequency Modes for Resting-State BOLD Activity based on MODWT and EMD

I estimated characteristic frequency per representative ROI time series per scan. For the MODWT decomposition scheme, I adopted the *pwelch* method as provided by MATLAB. For the EMD decomposition scheme, I adopted the *hht* method as provided by MATLAB. I first averaged the characteristic frequency per ROI across scans and afterward, I got the mean and standard deviation across the number of ROIs. Table 1 summarizes the whole-brain averaged characteristic intrinsic frequency modes for resting-state BOLD activity extracted with both filtering schemes from the MyConnectome dataset. The mean frequency of subbands between the two filtering methods doesn’t overlap. Similarly, I reported the whole-brain averaged characteristic intrinsic frequency modes for a resting-state BOLD activity for the NYU dataset averaging first across the brain per subject and scan, then averaging across scans, and finally estimating the mean and standard deviation across the cohort.

**Table 1.** Whole-brain averaged intrinsic frequency modes for both MODWT and EMD filtering schemes from the MyConnectome dataset.

|  | Subband 1 | Subband 2 | Subband 3 | Subband 4 |
| --- | --- | --- | --- | --- |
| **MODWT** | **0.21±0.004** | **0.11±0.004** | **0.049±0.004** | **0.021±0.004** |
| **EMD** | **0.27±0.012** | **0.14±0.011** | **0.072±0.006** | **0.023±0.003** |

### Researcher’s Free Choice Pre-processing Paths May Affect the Repeatability of Multi-Frequency Multilayer Network Topologies

Results of three-way ANOVA with repeated measures on three factors (filtering - connectivity estimator - topological layout) in the atlas and subnetworks space revealed an effect on the repeatability of multi-frequency multilayer network topologies (p < 0.05; corrected for multiple comparisons). Tables 3 and 4 tabulate the findings from the three-way ANOVA for the atlas and subnetworks level, correspondingly for the MyConnectome study. Similarly, STables 4-5 and STables 6-7 tabulate the reports of three-way ANOVA for the short-term and long-term repeat scan sessions, correspondingly for the NYU study. In simple words, the interpretation of the findings from the three ANOVA is that every choice across the three factors has a significant effect on the final multi-frequency multilayer network topology.

**Table 2.**
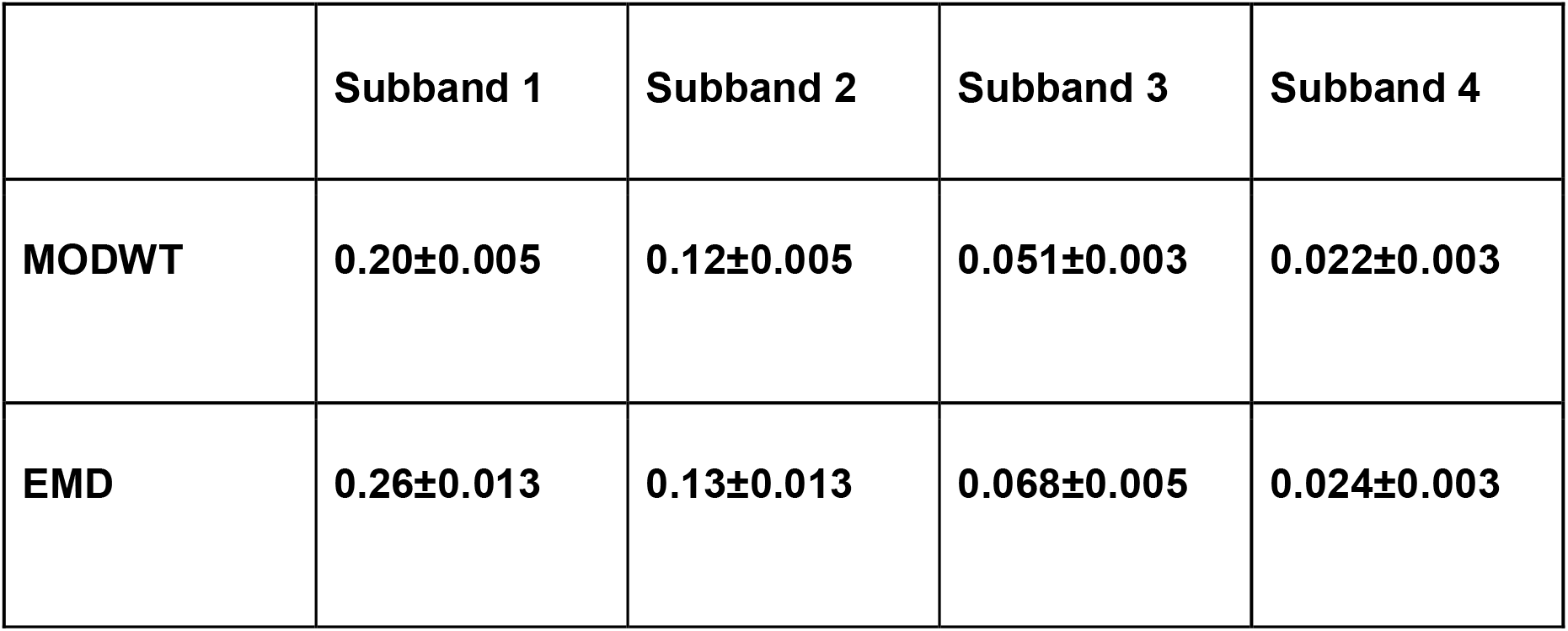
Whole-brain averaged intrinsic frequency modes for both MODWT and EMD filtering schemes from the NYU test-retest dataset.

**Table 3.**
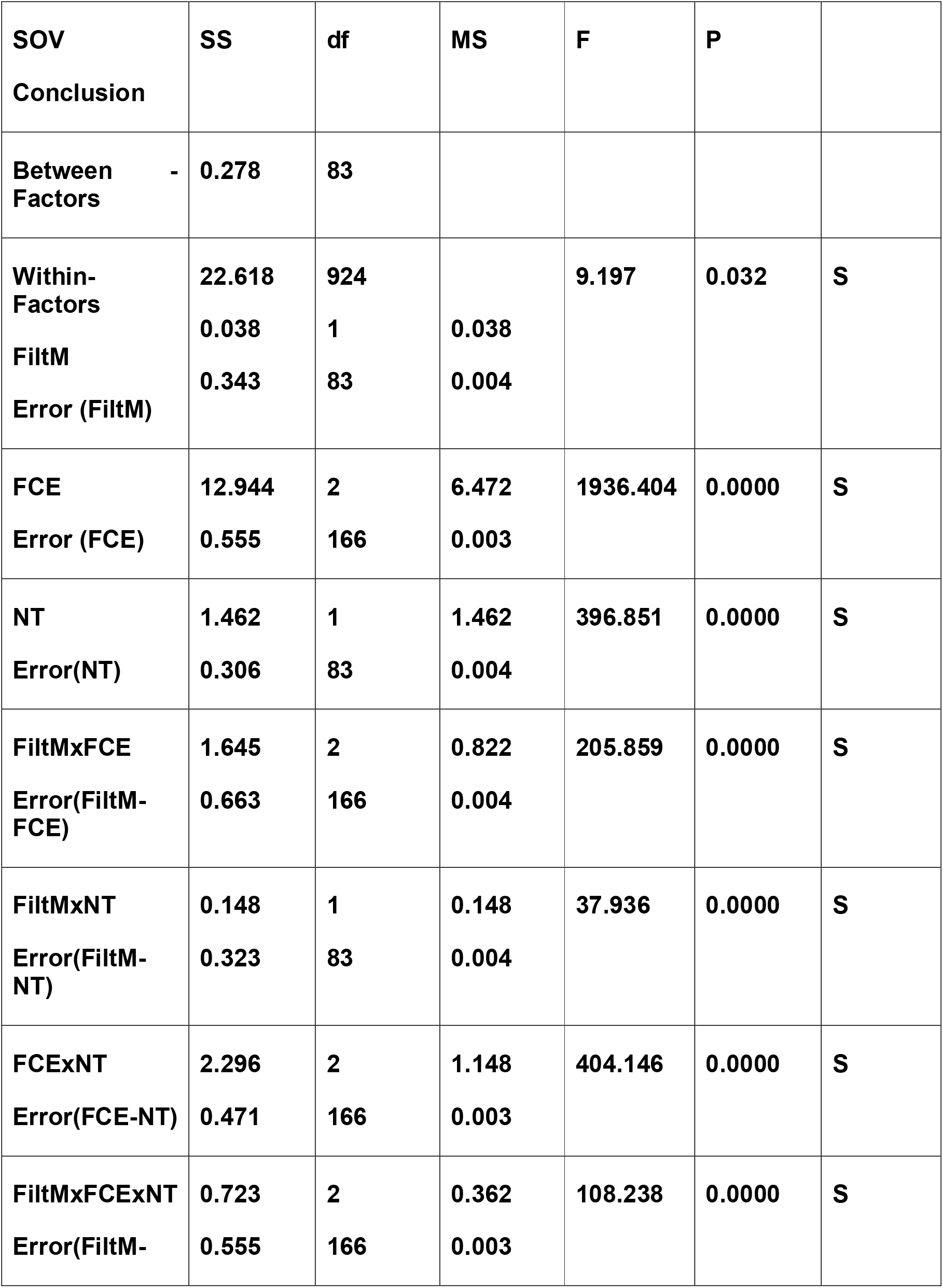
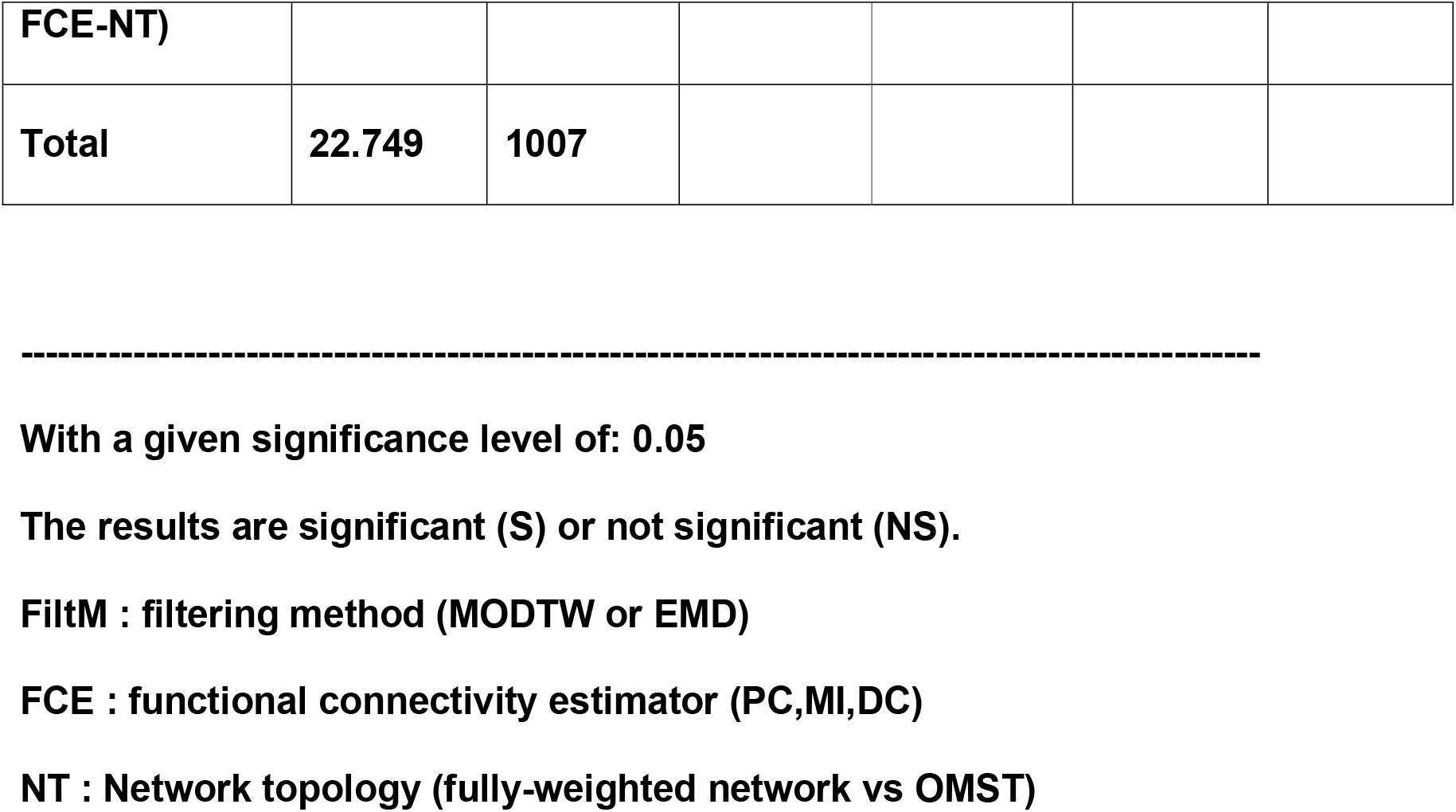
Three-Way Analysis of Variance With Repeated Measures on Three Factors (Within-Factors) based on Full Network Resolution Analysis for MyConnectome study (p < 0.05; corrected for multiple comparisons).

**Table 4.**
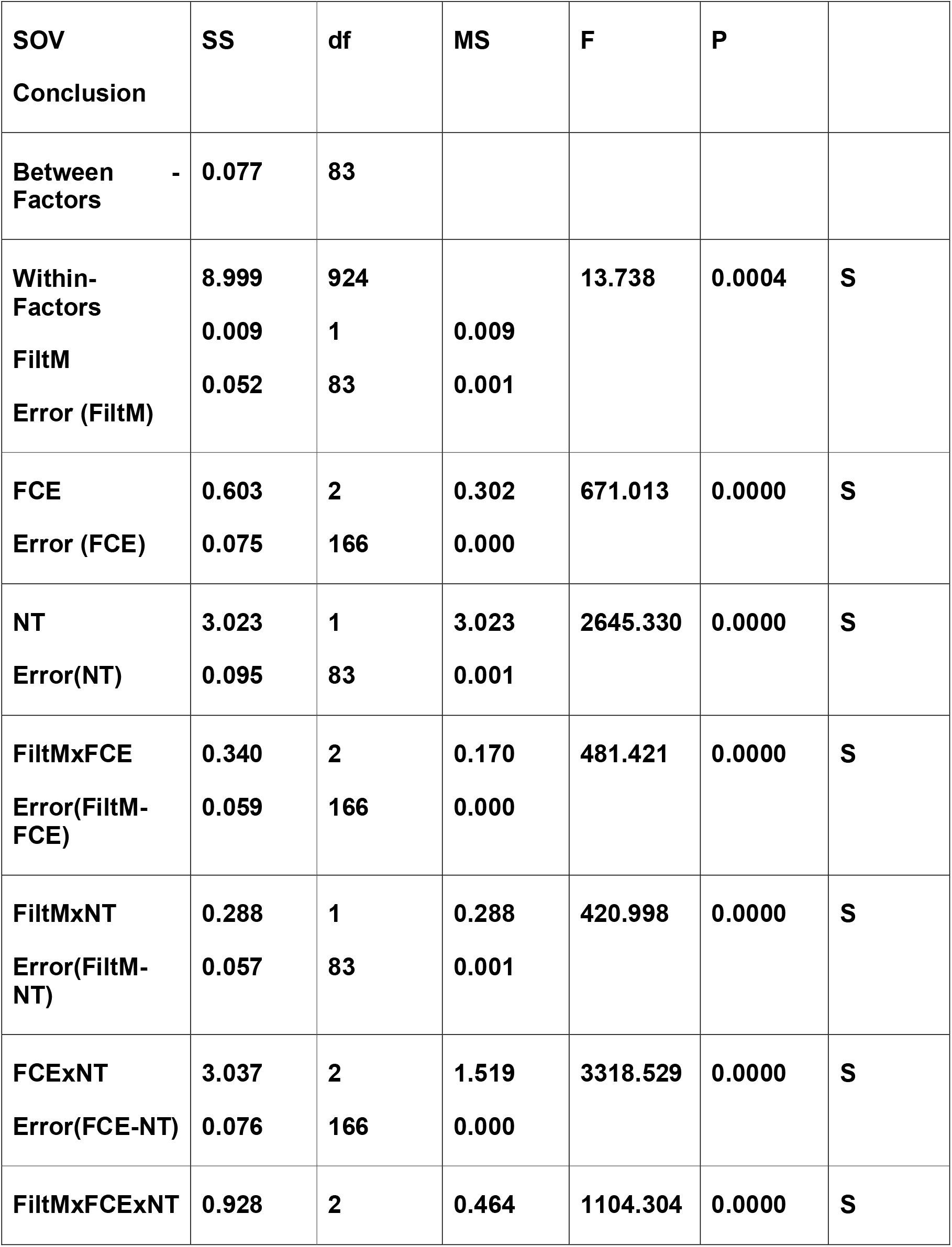
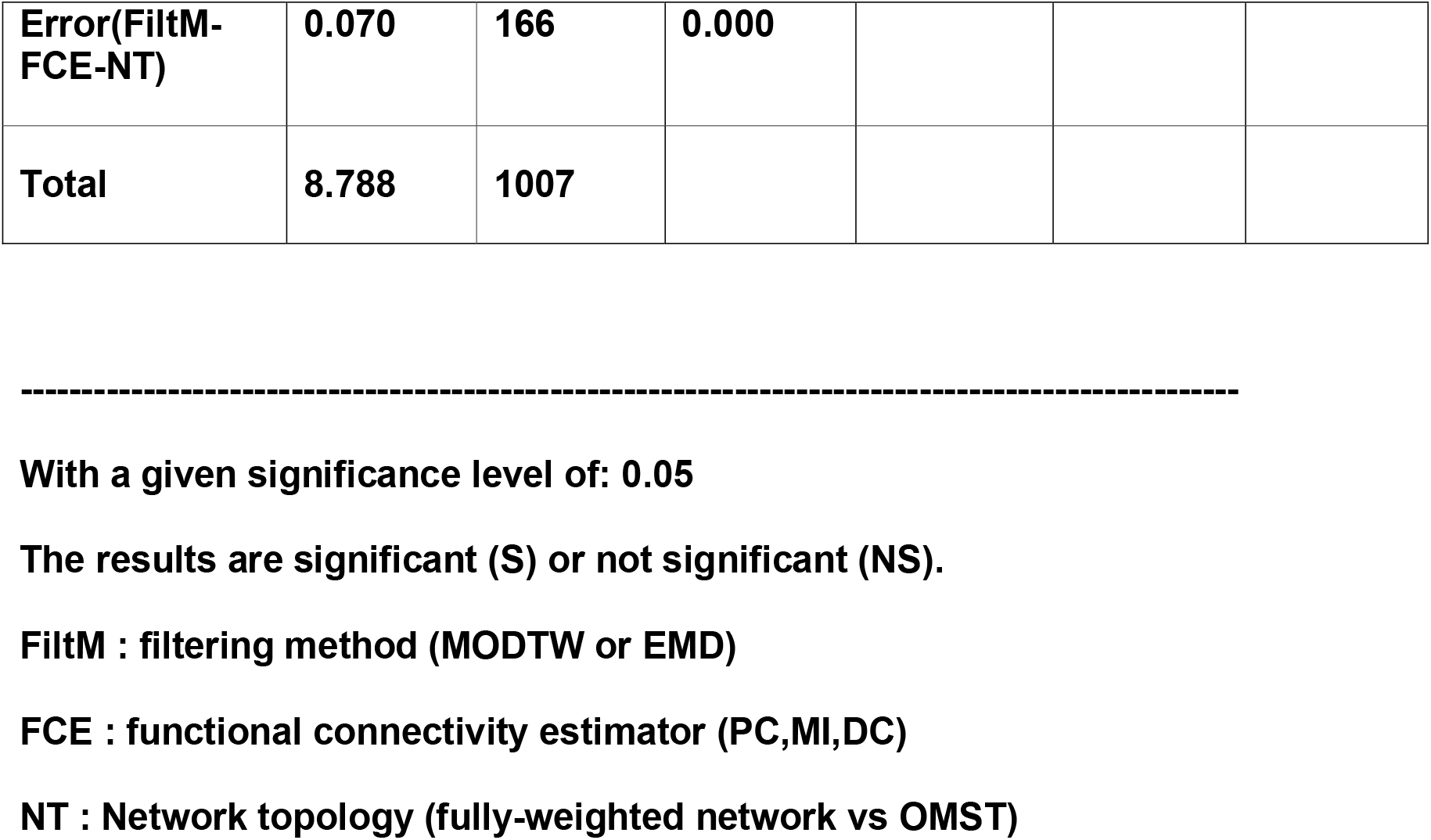
Three-Way Analysis of Variance With Repeated Measures on Three Factors (Within-Factors) based on Subnetwork Resolution Analysis for MyConnectome study (p < 0.05; corrected for multiple comparisons).

### PDiV reveals Important Topological Differences Across Pipelines

Fig.5 illustrates the across-scans PDiV values averaged and the relevant standard deviations per pipeline for the MyConnectome study. Figs 6 - 7 illustrate the across-subjects PDiV values and the relevant standard deviations per pipeline for the short-term and long-term repeat scan sessions for the NYU study. Below, I reported the repeatable pipelines for every case after applying the adopted statistical threshold of (*PDiV*_*p*_ ≤ *PDiV*^*threshold*^) :

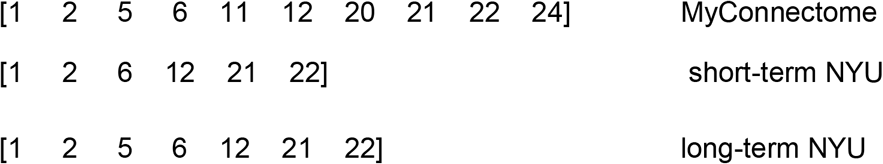

where 1 denotes the first left pipeline as reported in Figs. 5 - 7 (leftmost) while 24 is the last one (rightmost). The common pipelines across the three cases are the following six:

**Figure 5.**
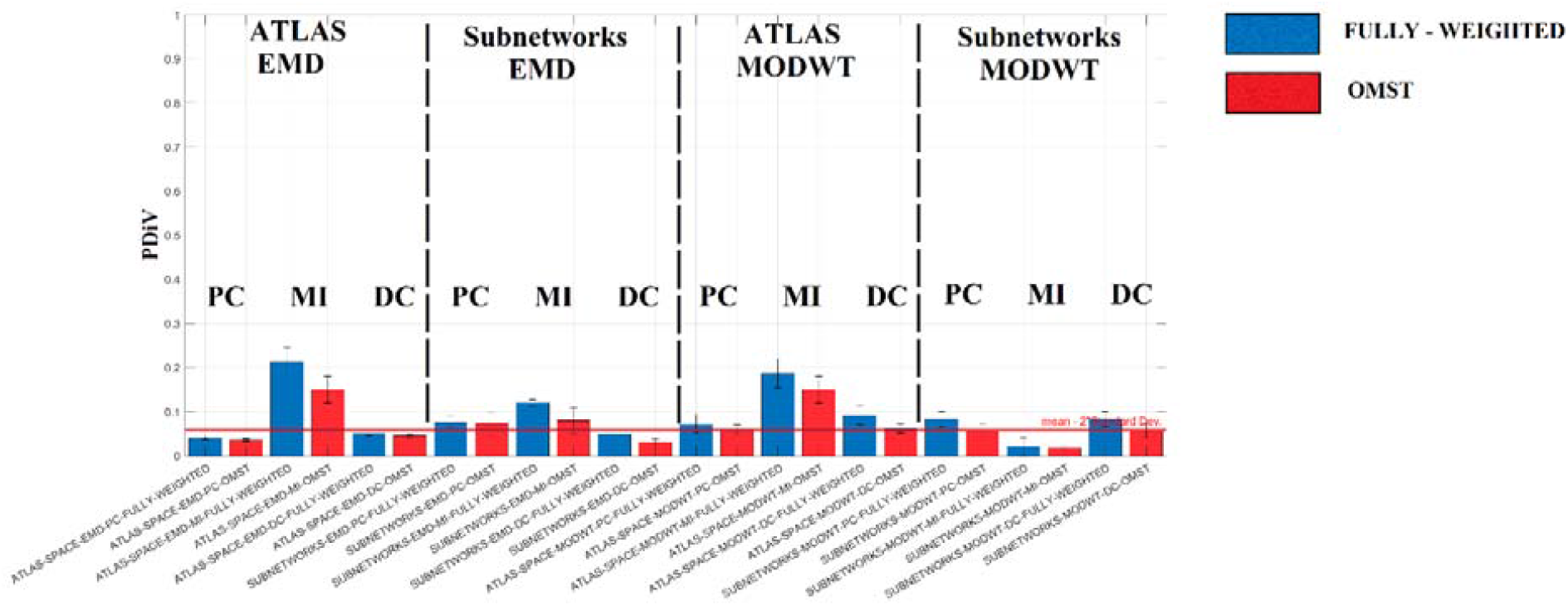
Repeatability of Multi-Frequency Multilayer Brain Network Topologies across distinct pipelines. PDiV scan-averaged values across every possible pipeline (24 in total) among the four factors explored in the MyConnectome dataset. The red line denotes the.

**Figure 6.**
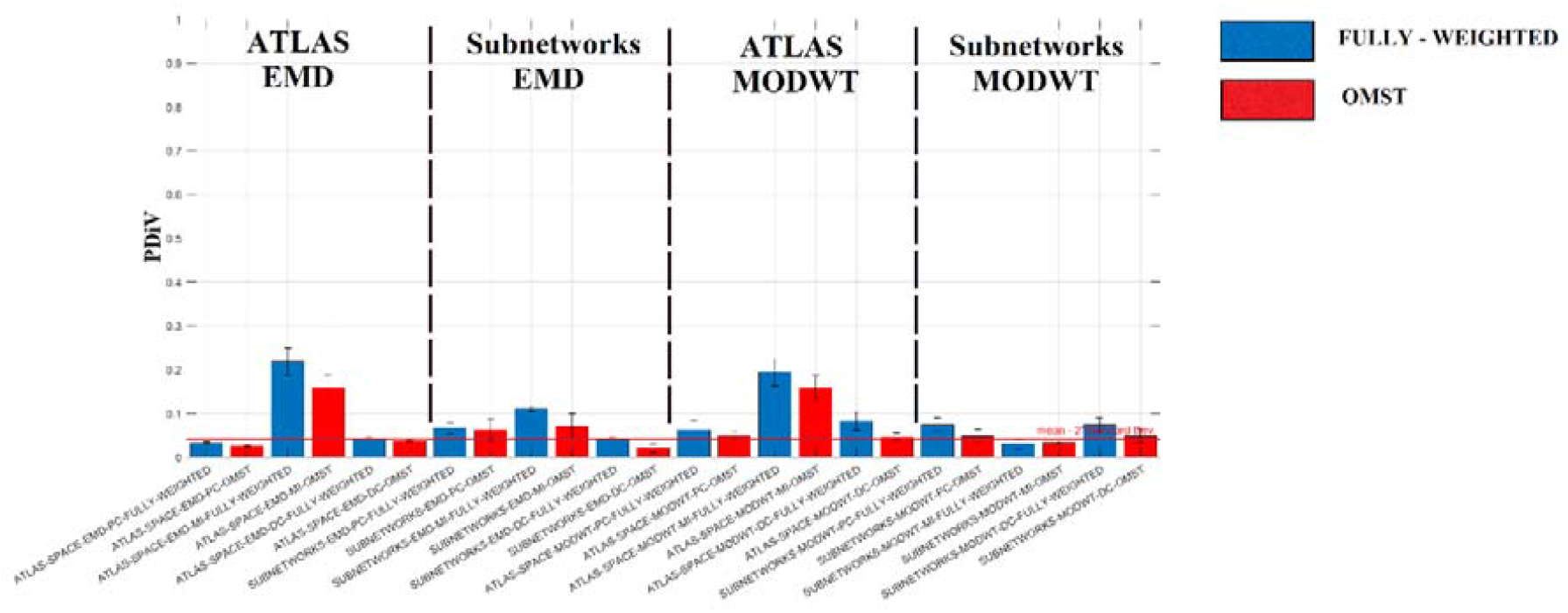
Repeatability of Multi-Frequency Multilayer Brain Network Topologies across distinct pipelines. PDiV scan-averaged values across every possible pipeline (24 in total) among the four factors explored in the short-term repeatability session of the NYU dataset. The red line denotes the.

1- ATLAS-SPACE-EMD-PC-FULLY-WEIGHTED

2- ATLAS-SPACE-EMD-PC-OMST

6 - ATLAS-SPACE-EMD-DC-OMST

12 - SUBNETWORKS-EMD-DC-OMST

21 - SUBNETWORKS-MODWT-MI-FULLY-WEIGHTED

22- SUBNETWORKS-MODWT-MI-OMST

### Evaluation of Pipelines based on the Within- vs Between-subject PDiV Criterion

My findings suggest that the proposed six pipelines based on the PDiV showed systematically a higher proportion of subjects with WS^PDiV^ values lower than BS^PDiV^ in both short-term and long-term time spans of the NYU dataset. Specifically, for the short-term time span, the average number of subjects with WS^PDiV^ values lower than BS^PDiV^ for the six suggested pipelines was 23 subjects and 13 for the rest of the pipelines (p = .000311). For the long-term time-span, the average number of subjects with WS^PDiV^ values lower than BS^PDiV^ for the six suggested pipelines was 22 subjects and 11 for the rest of the pipelines (p = .000432).

### Relationships between PDIVs and subject motion

For the NYU short-term and long-term datasets, the six proposed pipelines from the first criterion showed no significant correlation between PDiV values and subject motion. The rest of the 18 pipelines exhibited a significant correlation between PDiV value and motion with a magnitude ranging between 0.29 and 0.37 for the short-term dataset, and between 0.27 and 0.39 for the long-term dataset.

### Overall recommendations for adequate multi-frequency multilayer network construction pipelines

Based on the three adopted criteria, I final recommend the six pipelines as were detected from the first criterion and further evaluated by the other two. It is important to mention here that repeatability is preserved in both the atlas and subnetworks spatial layout. A significant outcome of my study is that EMD produces repeatable multi-frequency multilayer network topologies in both spatial scales, while MODWT is only for the subnetworks level. In four out of six pipelines, OMST data-driven topological filtering method is part of the best pipelines compared to two for the fully-weighted networks. For the adopted connectivity estimators, DC is the only estimator that participates in the best pipelines in both spatial scales but only in collaboration with EMD, PC is spatially connected to the atlas level and EMD filtering method, and MI is spatially connected to the subnetworks level and MODWT filtering method.

To further illustrate the (dis)similarity of multi-frequency multilayer topologies between the good and bad pipelines, I estimated their PDiV similarity in a pair-wise fashion, and then I projected these associations into a 3D space (see section 4 in supp.material). Interestingly, the good and bad pipelines form a cluster distant from each other that further support the whole analysis and the adopted criteria. The good pipelines produce similar topologies under the PDiV framework while simultaneously these network topologies are distant from the ones produced from the bad pipelines.

## DISCUSSION

A large amount of current neuroimaging research with fMRI is focused on harnessing repeatable brain network-based connectomic biomarkers related to both normal and abnormal brain function. However, this investigation involves a combination of arbitrary pre-processing choices (Korhonen et al. 2021). Test-retest repeatability is a prerequisite to the definition of repeatable connectomic biomarkers (Fornito et al. 2015; Hallquist and Hillary 2019). Here, I explored for the very first time in the literature how different researchers’ choices may affect the repeatability of multi-frequency multilayer network topologies. I systematically investigated 24 unique pipelines from resting-state fMRI recordings acquired from 84 scans of a single subject (MyConnectome dataset ;(Poldrack et al. 2015)) and 25 subjects with 3 repeat scan sessions in both short-term and long-term periods (NYU dataset; Shehzad et al., 2009). In the present study as in a previous one (Luppi et al., 2021), I adopted the PDiV metric as a proper way to evaluate the pipelines based on a multi-scale network topological comparison compared to adapting trivial network metrics. I also adopted three main criteria to uncover the pipelines that reproduce consistent brain network topologies. My findings suggest that the synergy of preprocessing steps produces repeatable brain network topologies and not just a single choice in any of them.

Test-retest studies of resting-state fMRI single-layer brain networks focused on the repeatability of graph metrics in various cohorts and both short and long-term periods between scans (Song et al. 2012; Andellini et al. 2015; Somandepalli et al. 2015; Shah et al. 2016; Termenon et al. 2016; Wang et al. 2017; Noble et al. 2017, 2019). However, the estimation of graph metrics derived from the network topology and the conclusion regarding the repeatability of these network metrics might differ from study to study due to the researcher’s choice of which to represent and also due to their different nature. For that reason, I adopted the PDiV metric as a proper measure to quantify brain network topology similarities in data acquired from the same subject across short (minutes), medium (weeks), and long (month) time periods in two open independent datasets.

In summary, my findings untangle large and medium systematic differences in the adopted pipelines in terms of constructing similar multi-frequency multilayer network topologies from two or more scans for the same subject. Importantly, my findings were consistent across short, medium, and long-term time-periods between scan sessions, in two independent rs-fMRI datasets. Remarkably that an inappropriate choice of the pipeline can impair the repeatability of network topology even for scans acquired less than 45 minutes apart up to a 10-fold increase in network topological dissimilarity (PDiV) compared to the best pipelines (Fig.6). The best performing pipelines can construct a repeatable multi-frequency multilayer network topology for the same subject, even at a time distance of many months (less than a year; Fig.7), and more than a year (Fig.5).

**Figure 7.**
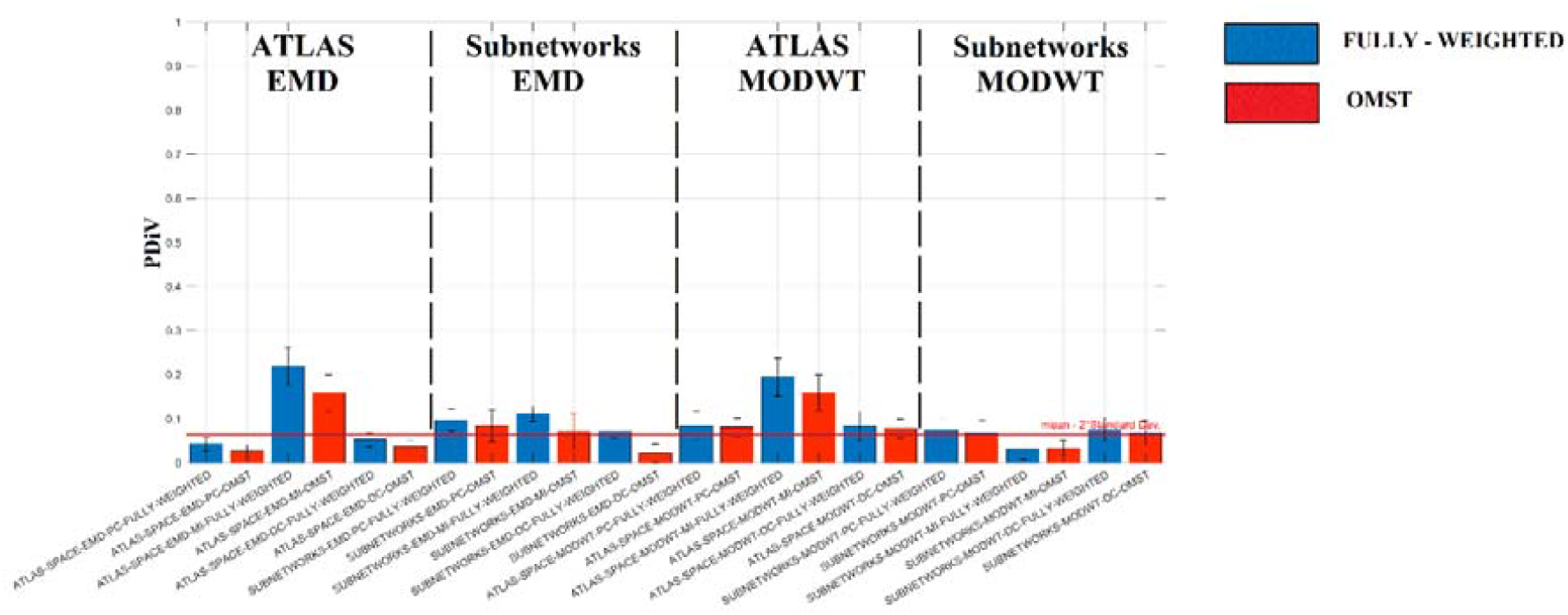
Repeatability of Multi-Frequency Multilayer Brain Network Topologies across distinct pipelines. PDiV scan-averaged values across every possible pipeline (24 in total) among the four factors explored in the long-term repeatability session of the NYU dataset. The red line denotes the.

These findings propose that an inappropriate choice of a multi-frequency multilayer network construction pipeline may have significant effects on brain network properties and conclusions in longitudinal studies (Zhang et al., 2021). Repeatability of multi-frequency multilayer network topologies from two scan sessions of the same subject is of paramount importance when a researcher wants to associate network properties with behavioral traits (Smith et al., 2013) or with clinical groups like schizophrenia, autism, or Alzheimer’s disease (Nichols et al., 2017; Gifford et al., 2020; Wei et al., 2022). Group comparison of network metrics derived from multi-frequency multilayer networks derived from magnetoencephalographic resting-state recordings has been also introduced (Mandke et al., 2018). The detection of repeatable multi-frequency multilayer network construction pipelines constitutes also a prerequisite for network exploratory analysis (Nichols et al., 2017) and clinical translation (Chen et al., 2018).

Hopefully, my analysis untangled six out of twenty-four multi-frequency multilayer network construction pipelines with consistent findings in two datasets (MyConnectome and NYU) and across different time spans. As I mentioned in the results section, I adopted MVPR as our unique algorithmic choice for the surrogate analysis and statistical topological filtering of the multi-frequency multilayer networks. My analysis showed repeatable network patterns to be preserved in both the atlas and subnetworks spatial layout. A significant outcome of our study is that EMD produces repeatable multi-frequency multilayer network topologies in both spatial scales, while MODWT is only for the subnetworks level. In four out of six pipelines, OMST data-driven topological filtering method is part of the best pipelines compared to two for the fully-weighted networks. For the adopted connectivity estimators, DC is the only estimator that participates in the best pipelines in both spatial scales but only in collaboration with EMD, while PC is spatially connected to the atlas level and EMD filtering method, and MI is spatially connected to the subnetworks level and MODWT filtering method.

In the present study, I decided to decompose the BOLD signal with a well-known technique, the MODWT (Zhang et al. 2016) vs an adaptive filtering technique called EMD (Yuen et al. 2019). I intended to demonstrate how different filtering techniques of the BOLD signal can lead to repeatable multi-frequency multilayer topologies as part of a common pre-processing pipeline. The mean frequency across the ROIs and scans of every frequency subband with both methods didn’t overlap. This finding could be evidence of the complementary information encapsulated by the ROI-based time series extracted from both filtering methods. This is a statement that deserves further consideration in a multi-subject test-retest study.

It is important to mention that repeatability is a significant criterion that neuroscientists should consider for their choice of the multi-frequency multilayer network construction pipeline. However, the adopted pipelines should also demonstrate to a large number of participants that the within-subjects (WS) PDiV is smaller than between-subjects (BS). In addition, the recommended pipelines should not show a significant correlation of the PDiV values with the subject’s motion (Luppi et al., 2021). The constructed networks should also be linked to neurobiologically supportive results (Shirer et al., 2015; Noble et al., 2019). Complementary to the aforementioned three criteria, it is significant to reveal network changes due to interventions (cognitive, physical training, both of them, and others), neuromodulations (task and arousal changes), pharmacological substances (caffeine, propofol, drugs, etc.), and behavioral history (cognition fatigue during the working day, sleep, etc). Current work should extend the validation of the proposed pipelines to detect associations of the multi-frequency multilayer network topologies to personality traits (Smith et al., 2015), and also to what extent they can increase the performance of a connectomic biomarker with the main aim to discriminate healthy individuals from various brain disorders (Chen et al., 2018).

### Limitations

In particular, I did not explore potential differences between resting-state conditions (eyes-open vs eyes-closed vs naturalistic viewing) (Van Dijk et al. 2010; Wang et al. 2011), or the impact of scan duration and arousal state (Laumann et al. 2017). Similarly, I did not consider a wide number of alternative parcellation schemes in existence but I adopted the parcellation scheme proposed by the authors provided for free in the MyConnectome dataset (Arslan et al. 2018; Eickhoff et al. 2018). Test-retest studies are the first step of a systematic evaluation of how alternative network processing steps can affect the repeatability of network topologies. Further investigation of the adequacy of the proposed pipelines should include their sensitivity to detect topological changes due to tasks, to pharmacological substances e.g. anesthesia (Luppi et al., 2021), and various brain diseases. We hope that the proposed network topology construction framework will lead to more consistent analytic practices in the human network neuroscience of functional neuroimaging data.

### Conclusions

In conclusion, my study provides an exploratory framework searching for the best multi-frequency multilayer network construction pipelines across 24 candidates, to recover a repeatable brain network topology. My findings support that only the combination of several specific processing steps can guarantee the repeatability of multi-frequency multilayer network topologies. I untangled that every choice across the adopted processing steps matters and specific pipelines can produce similar network topologies over time-periods that span from minutes to several months evaluated in two independent datasets. Interestingly, alternative pipelines produce repeatable multi-frequency multilayer networks leading to the assumption that they share complementary information.

## Acknowledgements

SID is supported by a Beatriu de Pinós fellowship (2020 BP 00116). SID was supported by MRC grant <u>MR/K004360/1</u> and a Marie Sklodowska-Curie COFUND EU-UK Research Fellowship.

## Preprint

This study has been submitted to biorxiv as a preprint research article and can be found at the following link: https://www.biorxiv.org/content/10.1101/2021.10.10.463799v1

## Data and code availability

The NYU dataset is freely available from the International Neuroimaging Data-Sharing Initiative (INDI) (http://www.nitrc.org/projects/nyu_trt). Python code for the portrait divergence is freely available online (https://github.com/bagrow/network-portrait-divergence). MATLAB code for the Orthogonal Minimum Spanning Tree thresholding is freely available online (https://github.com/stdimitr/topological_filtering_networks).

