## Supplementary Material for "Assessing The Repeatability of Multi-Frequency Multi-Layer Brain Network Topologies Across Alternative Researcher’s Choice Paths"

Stavros I. Dimitriadis^1-8^*

^1^Department of Clinical Psychology and Psychobiology, Faculty of Psychology, University of Barcelona, Passeig de la Vall d’Hebron, 171, 08035, Barcelona, Spain

^2^Institut de Neurociències, University of Barcelona, Campus Mundet, Edifici de Ponent , Passeig de la Vall d’Hebron, 171, 08035 , Barcelona, Spain

^3^Integrative Neuroimaging Lab, 55133, Thessaloniki, Greece

^4^Cardiff University Brain Research Imaging Centre (CUBRIC), School of Psychology, College of Biomedical and Life Sciences, Cardiff University, CF24 4HQ, Cardiff, Wales, United Kingdom

^5^Neuroinformatics Group, Cardiff University Brain Research Imaging Centre (CUBRIC), School of Psychology, College of Biomedical and Life Sciences, CF24 4HQ, Cardiff, Wales, United Kingdom

^6^Division of Psychological Medicine and Clinical Neurosciences, School of Medicine, College of Biomedical and Life Sciences, Cardiff University, CF24 4HQ, Cardiff, Wales, United Kingdom

^7^Neuroscience and Mental Health Research Institute, School of Medicine, College of Biomedical and Life Sciences, Cardiff University, CF24 4HQ, Cardiff, Wales, United Kingdom

^8^MRC Centre for Neuropsychiatric Genetics and Genomics, School of Medicine, College of Biomedical and Life Sciences, Cardiff University, CF24 4HQ, Cardiff, Wales, United Kingdom

* corresponding author: Dr.Stavros I. Dimitriadis

1. **Date preprocessing:**

All fMRI data were preprocessed according to a pipeline developed at Washington University (Poldrack et al., 2015). First, the initial 100 time points were discarded from the data because of the presence of an evoked auditory signal within the MRI scanner. Data were realigned to correct for head motion and normalized to a mode of 1,000. Each session was registered to a single session that had previously been registered to the mean T1-weighted structural image and atlas. The session to atlas transform was inverted and applied to the mean field map so that the distortion correction could be applied in each session’s space. The undistorted data were then reregistered to the atlas space. The transforms for head motion correction and affine registration to atlas space were combined with the field map-based distortion correction to resample the data from the original session space to the undistorted 3-mm isotropic atlas space in a single step using FSLs applywarp tool.

Artifacts were reduced using frame censoring, regression, and spectral filtering. Frames with framewise displacement >0.25 mm were censored as well as uncensored segments of data lasting less than five contiguous volumes (97.1 ± 4% of frames were kept). Nuisance regressors included whole brain, white matter, and ventricular signals and their derivatives in addition to 24 movement regressors derived by Volterra expansion. Interpolation over censored frames was computed by least squares spectral estimation so that continuous data could be band pass-filtered (0.01–0.1 Hz); 12 of 104 sessions were discarded after an alteration to the scanning protocol, and 8 were discarded because of poor normalization.

**Parcellation Scheme.**

After preprocessing, the mean time series was extracted from 630 predefined regions of interest (ROIs): 616 cortical parcels (309 from the left hemisphere and 307 from the right hemisphere) from a predefined cortical parcellation scheme (Laumann et al., 2015) that was based on the principles of the Gordon atlas (Poldrack et al., 2015) but used all 84 sessions of the individuals’ data to create a more subject-specific parcellation scheme along with 14 subcortical parcels from the Harvard–Oxford subcortical atlas ([fsl.fmrib.ox.ac.uk/fsl/fslwiki/](http://fsl.fmrib.ox.ac.uk/fsl/fslwiki/)).

2. **Evaluation of Surrogate Models**

SFigure 1 illustrates how the adapted five properties were computed for every ROI-based time-series and the relevant surrogates produced by the two surrogate null-models. STables 1 – 3 tabulate the percentage of ROIs that showed significant difference of at least three properties between the two surrogate null-models. I practically compared the aPCC – MI values estimated for the five properties for every ROI, filtering method, and subbands between the original and surrogate time-series. We finally reported the scan-based (MyConnectome study) and group-based (NYU study) percentage of ROIs that demonstrated differences to at least three properties between the two surrogate null models (STables 1 - 3). MVPR method showed significant lower aPCC – MI values compared to MVAR method which practically means that the surrogate time-series mimic better the original time-series.

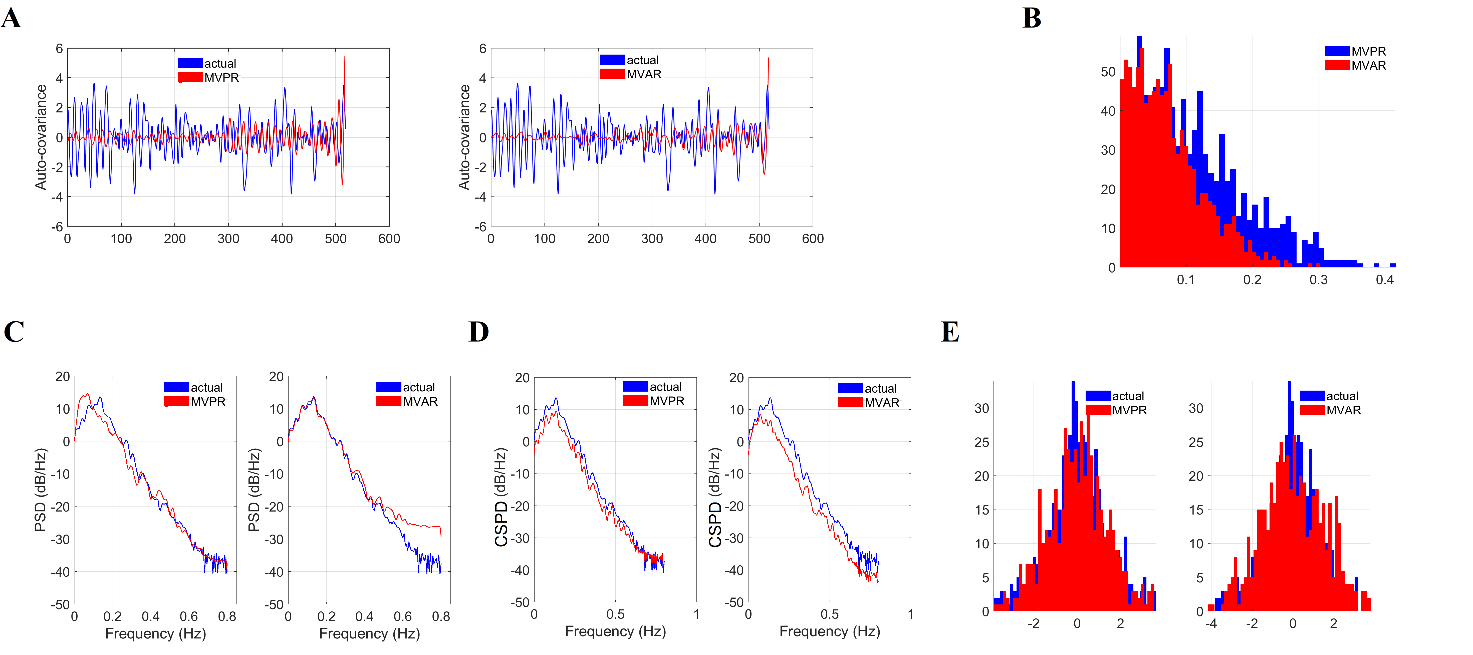

**SFigure 1.** Demonstration of how the adapted properties were computed for an example of an ROI-based representative time-series and each surrogate produced with both surrogate null models.

A. the auto-covariance of the original time-series with the surrogates produced with both models

B. the distribution of the absolute value of Pearson’s correlation coefficient (aPCC) between the original time-series and 1.000 surrogates produced with both models

C. the power-spectral density of the original time-series with the surrogates produced with both models

D. the cross-power spectral density between t the original time-series with surrogates produced with both models

E. the distributions of the amplitude of the original time-series and the surrogates in a common plot

**MyConnectome Study:**

**STable 1.** Percentage of ROIs that shows a significant difference between the two surrogate null-models to at least 3 properties. My findings are reported in every sub-band and in both filtering methods.

|  | **Subband 1** | **Subband 2** | **Subband 3** | **Subband 4** |
| --- | --- | --- | --- | --- |
| **EMD** | 29.37 ± 3.76 % | 26.19 ± 4.12 % | 27.62 ± 4.91 % | 28.73 ± 3.07 % |
| **MODWT** | 26.19 ± 3.54 % | 27.14 ± 5.42 % | 28.89 ± 4.28 % | 30.32 ± 3.88 % |

**NYU Study:**

**STable 2 (short-term).** Percentage of ROIs that shows a significant difference between the two surrogate null-models to at least 3 properties. My findings are reported in every sub-band and in both filtering methods. Mean and standard deviation were estimated across the subjects

|  | **Subband 1** | **Subband 2** | **Subband 3** | **Subband 4** |
| --- | --- | --- | --- | --- |
| **EMD** | 32.13 ± 3.21 % | 30.65 ± 1.98 % | 33.51 ± 3.32 % | 32.82 ± 3.22 % |
| **MODWT** | 31.42 ± 2.65 % | 29.44 ± 2.32 % | 32.72 ± 4.12 % | 34.54 ± 2.92 % |

**STable 3 (long-term).** Percentage of ROIs that shows a significant difference between the two surrogate null-models to at least 3 properties. My findings are reported in every sub-band and in both filtering methods. Mean and standard deviation were estimated across the subjects

|  | **Subband 1** | **Subband 2** | **Subband 3** | **Subband 4** |
| --- | --- | --- | --- | --- |
| **EMD** | 35.65 ± 4.12 % | 34.77 ± 2.25 % | 32.48 ± 3.12 % | 36.72 ± 3.43 % |
| **MODWT** | 32.51 ± 3.23 % | 35.18 ± 2.97 % | 35.67 ± 4.56 % | 35.81 ± 3.88 % |

3. **Three – way ANOVA on Short – term and long-term repeat-scan sessions of NYU study**

Results of three-way ANOVA with repeated measures on three factors (filtering - connectivity estimator - topological layout) in the atlas and subnetworks space revealed an effect on the repeatability of multi-frequency multilayer network topologies (p < 0.05; corrected for multiple comparisons). STables 4-5 and STables 6-7 tabulate the reports of three-way ANOVA for the short-term and long-term repeat scan sessions, correspondingly for the NYU study.

**STable1.** Three-Way Analysis of Variance With Repeated Measures on Three Factors (Within-Factors) based on Full Network Resolution Analysis for the short-term NYU dataset (p < 0.05; corrected for multiple comparisons).

| **SOV**  **Conclusion** | **SS** | **df** | **MS** | **F** | **P** |  |
| --- | --- | --- | --- | --- | --- | --- |
| **Between -Factors** | **0.155** | **24** |  |  |  |  |
| **Within-Factors**  **FiltM**  **Error (FiltM)** | **8.937**  **0.018**  **0.228** | **275**  **1**  **24** | **0.008**  **0.001** | **2.880** | **0.018** | **S** |
| **FCE**  **Error (FCE)** | **4.745**  **0.307** | **2**  **48** | **2.372**  **0.006** | **370.497** | **0.0000** | **S** |
| **NT**  **Error(NT)** | **0.561**  **0.170** | **1**  **24** | **0.561**  **0.007** | **79.280** | **0.0000** | **S** |
| **FiltMxFCE**  **Error(FiltM-FCE)** | **0.483**  **0.421** | **2**  **48** | **0.241**  **0.009** | **27.549** | **0.0000** | **S** |
| **FiltMxNT**  **Error(FiltM-NT)** | **0.046**  **0.202** | **1**  **24** | **0.046**  **0.008** | **5.433** | **0.0000** | **S** |
| **FCExNT**  **Error(FCE-NT)** | **0.803**  **0.305** | **2**  **48** | **0.401**  **0.006** | **63.150** | **0.0000** | **S** |
| **FiltMxFCExNT**  **Error(FiltM-FCE-NT)** | **0.272**  **0.332** | **2**  **48** | **0.136**  **0.007** | **19.614** | **0.0000** | **S** |
| **Total** | **9.047** | **299** |  |  |  |  |

**---------------------------------------------------------------------------------------------------**

**With a given significance level of: 0.05**

**The results are significant (S) or not significant (NS).**

**FiltM : filtering method (MODTW or EMD)**

**FCE : functional connectivity estimator (PC,MI,DC)**

**NT : Network topology (fully-weighted network vs OMST)**

**STable 2.** Three-Way Analysis of Variance With Repeated Measures on Three Factors (Within-Factors) based on Subnetwork Resolution Analysis for the short-term NYU dataset (p < 0.05; corrected for multiple comparisons).

| **SOV**  **Conclusion** | **SS** | **df** | **MS** | **F** | **P** |  |
| --- | --- | --- | --- | --- | --- | --- |
| **Between -Factors** | **0.038** | **24** |  |  |  |  |
| **Within-Factors**  **FiltM**  **Error (FiltM)** | **2.732**  **0.007**  **0.032** | **275**  **1**  **24** | **0.007**  **0.001** | **5.600** | **0.0264** | **S** |
| **FCE**  **Error (FCE)** | **0.164**  **0.031** | **2**  **148** | **0.082**  **0.001** | **128.717** | **0.0000** | **S** |
| **NT**  **Error(NT)** | **0.883**  **0.042** | **1**  **24** | **0.883**  **0.002** | **507.319** | **0.0000** | **S** |
| **FiltMxFCE**  **Error(FiltM-FCE)** | **0.090**  **0.025** | **2**  **48** | **0.045**  **0.001** | **85.827** | **0.0000** | **S** |
| **FiltMxNT**  **Error(FiltM-NT)** | **0.111**  **0.022** | **1**  **24** | **0.111**  **0.001** | **122.891** | **0.0000** | **S** |
| **FCExNT**  **Error(FCE-NT)** | **0.888**  **0.029** | **2**  **48** | **0.444**  **0.001** | **727.235** | **0.0000** | **S** |
| **FiltMxFCExNT**  **Error(FiltM-FCE-NT)** | **0.269**  **0.030** | **2**  **48** | **0.134**  **0.001** | **216.505** | **0.0000** | **S** |
| **Total** | **2.658** | **299** |  |  |  |  |

**---------------------------------------------------------------------------------------------------**

**With a given significance level of: 0.05**

**The results are significant (S) or not significant (NS).**

**FiltM : filtering method (MODTW or EMD)**

**FCE : functional connectivity estimator (PC,MI,DC)**

**NT : Network topology (fully-weighted network vs OMST)**

**STable 3.** Three-Way Analysis of Variance With Repeated Measures on Three Factors (Within-Factors) based on Full Network Resolution Analysis for the long-term NYU dataset (p < 0.05; corrected for multiple comparisons).

| **SOV**  **Conclusion** | **SS** | **df** | **MS** | **F** | **P** |  |
| --- | --- | --- | --- | --- | --- | --- |
| **Between -Factors** | **0.05** | **24** |  |  |  |  |
| **Within-Factors**  **FiltM**  **Error (FiltM)** | **8.168**  **0.004**  **0.056** | **275**  **1**  **24** | **0.004**  **0.002** | **2.899** | **0.0181** | **S** |
| **FCE**  **Error (FCE)** | **3.743**  **0.068** | **2**  **48** | **1.872**  **0.001** | **1315.774** | **0.0000** | **S** |
| **NT**  **Error(NT)** | **0.446**  **0.057** | **1**  **24** | **0.446**  **0.002** | **186.283** | **0.0000** | **S** |
| **FiltMxFCE**  **Error(FiltM-FCE)** | **0.447**  **0.113** | **2**  **48** | **0.224**  **0.002** | **94.629** | **0.0000** | **S** |
| **FiltMxNT**  **Error(FiltM-NT)** | **0.041**  **0.057** | **1**  **24** | **0.041**  **0.002** | **17.246** | **0.0004** | **S** |
| **FCExNT**  **Error(FCE-NT)** | **0.726**  **0.065** | **2**  **48** | **0.098**  **0.001** | **268.659** | **0.0000** | **S** |
| **FiltMxFCExNT**  **Error(FiltM-FCE-NT)** | **0.196**  **0.108** | **2**  **48** | **0.098**  **0.002** | **43.610** | **0.0000** | **S** |
| **Total** | **6.177** | **299** |  |  |  |  |

**---------------------------------------------------------------------------------------------------**

**With a given significance level of: 0.05**

**The results are significant (S) or not significant (NS).**

**FiltM : filtering method (MODTW or EMD)**

**FCE : functional connectivity estimator (PC,MI,DC)**

**NT : Network topology (fully-weighted network vs OMST)**

**STable 4.** Three-Way Analysis of Variance With Repeated Measures on Three Factors (Within-Factors) based on Subnetwork Resolution Analysis for the long-term NYU dataset (p < 0.05; corrected for multiple comparisons).

| **SOV**  **Conclusion** | **SS** | **df** | **MS** | **F** | **P** |  |
| --- | --- | --- | --- | --- | --- | --- |
| **Between -Factors** | **0.025** | **24** |  |  |  |  |
| **Within-Factors**  **FiltM**  **Error (FiltM)** | **6.535**  **0.001**  **0.010** | **275**  **1**  **24** | **0.001**  **0.000** | **5.127** | **0.0008** | **S** |
| **FCE**  **Error (FCE)** | **0.159**  **0.020** | **2**  **48** | **0.080**  **0.000** | **193.854** | **0.0000** | **S** |
| **NT**  **Error(NT)** | **0.902**  **0.034** | **1**  **24** | **0.902**  **0.001** | **628.412** | **0.0000** | **S** |
| **FiltMxFCE**  **Error(FiltM-FCE)** | **0.090**  **0.013** | **2**  **48** | **0.045**  **0.000** | **167.035** | **0.0000** | **S** |
| **FiltMxNT**  **Error(FiltM-NT)** | **0.076**  **0.016** | **1**  **24** | **0.076**  **0.001** | **113.679** | **0.0000** | **S** |
| **FCExNT**  **Error(FCE-NT)** | **0.843**  **0.021** | **2**  **48** | **0.422**  **0.000** | **973.526** | **0.0000** | **S** |
| **FiltMxFCExNT**  **Error(FiltM-FCE-NT)** | **0.255**  **0.017** | **2**  **48** | **0.128**  **0.000** | **363.412** | **0.0000** | **S** |
| **Total** | **2.483** | **299** |  |  |  |  |

**---------------------------------------------------------------------------------------------------**

**With a given significance level of: 0.05**

**The results are significant (S) or not significant (NS).**

**FiltM : filtering method (MODTW or EMD)**

**FCE : functional connectivity estimator (PC,MI,DC)**

**NT : Network topology (fully-weighted network vs OMST)**

4. **(Dis)similarity of good and bad pipelines under PDiV**

I constructed a topological distance PDiV matrix between every pair of pipelines and then we projected it into a 3D space. I adopted multidimensional scaling to project the PDiV matrix. I followed the same procedure for every dataset revealing an important trend : the good and bad pipelines form clusters that are separated from each other in the common 3D space. This practically means that the good pipelines produce similar multi-frequency multilayer topologies while simultaneously these patterns are distinct topologically from the topologies produced from the bad pipelines.

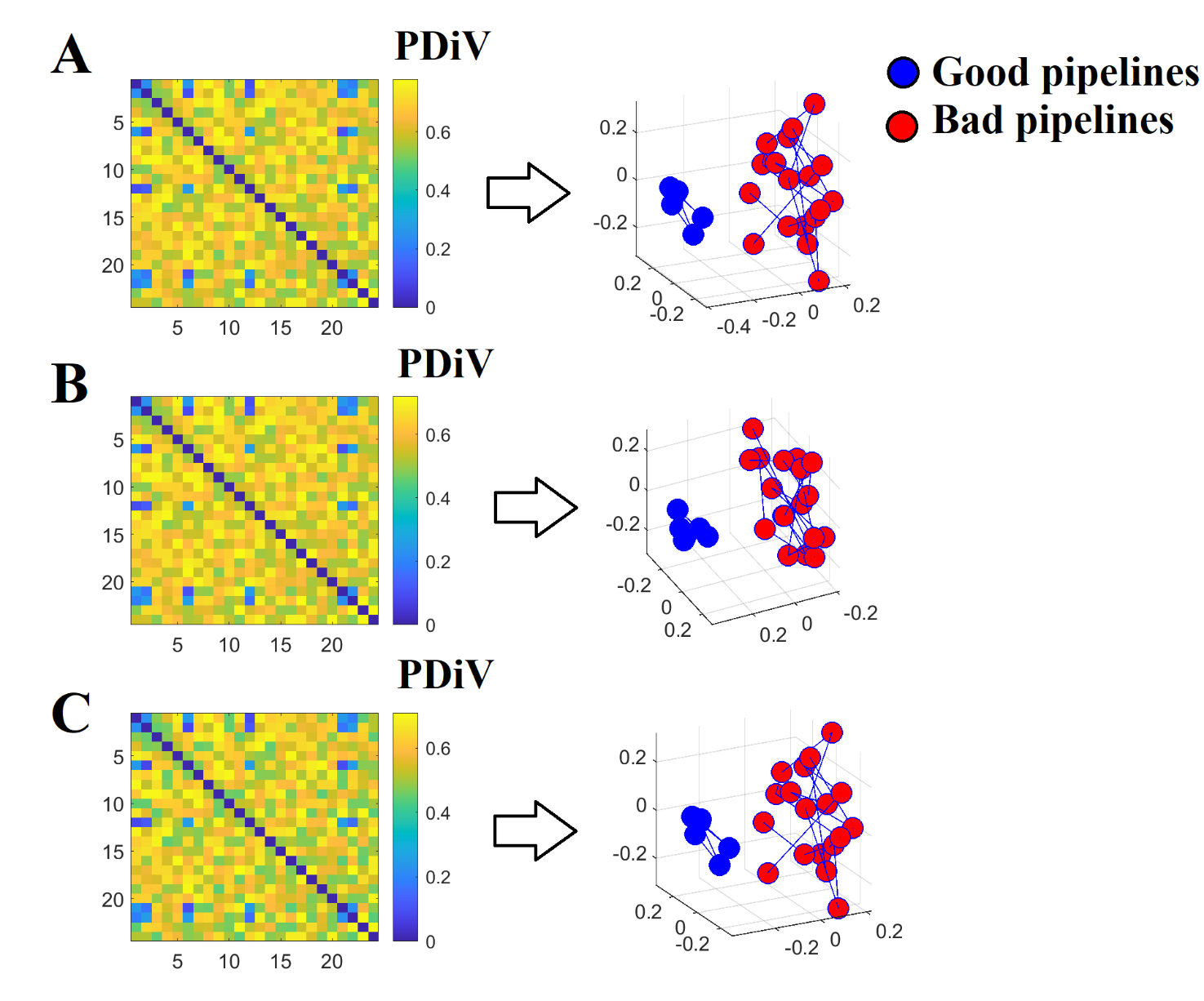

**SFigure 2.** PDiV (dis)similarity matrices between every pair of pipelines and the related projection into a 3D space. I projected every PDiV matrix with the multidimensional scaling algorithm. The stress function was < 0.1 which means that the whole procedure preserves the associations between the pipelines. Interestingly, the good and bad pipelines form a cluster separated from each other.

A) PDiV distance matrix between every pair of pipelines averaged across the 84 scans from the MyConnectome study
B) PDiV distance matrix between every pair of pipelines averaged across participants from NYU short-term study
C) PDiV distance matrix between every pair of pipelines averaged across participants from NYU long-term study
